# Hierarchical organization of spontaneous co-fluctuations in densely-sampled individuals using fMRI

**DOI:** 10.1101/2022.03.06.483045

**Authors:** Richard F. Betzel, Sarah A. Cutts, Jacob Tanner, Sarah A. Greenwell, Thomas Varley, Joshua Faskowitz, Olaf Sporns

## Abstract

Edge time series decompose FC into its framewise contributions. Previous studies have focused on characterizing the properties of high-amplitude frames, including their cluster structure. Less is known about middle- and low-amplitude co-fluctuations. Here, we address those questions directly, using data from two dense-sampling studies: the MyConnectome project and Midnight Scan Club. We develop a hierarchical clustering algorithm to group peak co-fluctuations of all magnitudes into nested and multi-scale clusters based on their pairwise concordance. At a coarse scale, we find evidence of three large clusters that, collectively, engage virtually all canonical brain systems. At finer scales, however, each cluster is dissolved, giving way to increasingly refined patterns of co-fluctuations involving specific sets of brain systems. We also find an increase in global co-fluctuation magnitude with hierarchical scale. Finally, we comment on the amount of data needed to estimate co-fluctuation pattern clusters and implications for brain-behavior studies. Collectively, the findings reported here fill several gaps in current knowledge concerning the heterogeneity and richness of co-fluctuation patterns as estimated with edge time series while providing some practical guidance for future studies.

## INTRODUCTION

The human brain can be modeled as a network of functionally interconnected brain regions [1]. In many applications, the weights of functional connections are defined as the bivariate correlation between two regions’ activity time series. Strong correlations are generally treated as evidence of functional connectivity (FC) [2].

Recent work has demonstrated that a static correlation between two time series, i.e. a functional connection, can be “temporally unwrapped” and precisely decomposed into its time-varying contributions [3–5]. This procedure generates a co-fluctuation or “edge time series”, whose elements indicate the magnitude and direction of instantaneous coupling between pairs of regions.

Previous analyses of edge time series have focused on high-amplitude co-fluctuations or”events” (but see [5] for an exception). Although events occur briefly and infrequently, the pattern of whole-brain co-fluctuations expressed during these periods necessarily contribute more to the time-averaged FC than lower-amplitude frames [3]. Moreover, high-amplitude co-fluctuation patterns can be partitioned into a small number of recurring clusters or “states” [6–8], encode information about subjects’ brain-based fingerprints [9], and can possibly enhance brain-behavior correlations [3].

However, interest in high-amplitude events – including work that predates our own [10–14] – has come at the expense of lower-amplitude co-fluctuations. In fact, very little is known about their properties. For instance, do low-amplitude frames exhibit cluster structure? Do they contain subject-specific information? How much does the inclusion of lower-amplitude peaks improve predictions of time-averaged FC?

Here, we address those questions directly. Specifically, we analyze densely-sampled data from the MyConnectome Project [15, 16] and Midnight Scan Club (MSC; five hours of data for ten subjects) [17, 18]. Focusing on peaks of the global co-fluctuation signal, we sample patterns corresponding to different magnitudes and cluster them using a bespoke hierarchical clustering algorithm. We discover that, while high- and low-amplitude co-fluctuation patterns form inter-mixed clusters, lower-amplitude patterns tend to be dissimilar from all other patterns and therefore less likely to participate in cohesive clusters. We investigate the hierarchical clusters in greater detail and show that, at a coarse scale, the majority of co-fluctuation patterns could be explained by three broad clusters that get sub-divided and refined at deeper hierarchical levels. Whereas the coarse clusters disclose broad, brain-wide co-fluctuation patterns, finescale clusters emphasize co-fluctuations involving distinct combinations of functional systems/networks. Finally, we reveal that accurately estimating cluster centroids requires large amounts of data and that, while coarse clusters “lock in” a basic pattern of FC, predictions of FC benefit from the inclusion of finescale clusters. This work is the first to investigate the organization of sub-event co-fluctuations, revealing rich structure while setting the stage for future studies.

## RESULTS

Here, we aimed to characterize co-fluctuation patterns estimated using “edge time series”. Briefly, this procedure entails z-scoring parcel time series, generating edge time series for every pair of parcels, and calculating the root mean square (RMS) of co-activity at each time point. We elected to focus on local maxima in this RMS time series – “peaks” – rather than all frames, as our previous studies using this same dataset demonstrated that “troughs” in the RMS signal correspond to highly variable co-fluctuation patterns [7]. After motion censoring, we detected a total of 3124 peaks. We further discarded peaks whose prominence (height minus the largest of its temporally adjacent troughs) was less than a value of 0.25 or occurred within 10 seconds of another peak. This procedure resulted in 1568 co-fluctuation patterns (whose statistical properties are described in Fig. S1; we show analogous statistics for data from the Midnight Scan Club in Fig. S2). For all subsequent analyses, we pooled the corresponding co-fluctuation patterns from scans.

### Cluster structure of full-spectrum co-fluctuation peaks

Previous studies demonstrated that high-amplitude co-fluctuations could be clustered into a small set of patterns, each of which recurred over time. However, those studies discarded all but highest-amplitude frames, i.e. putative “events”, and used a clustering algorithm that generated communities corresponding to a single organizational scale. Here, we extend those studies by clustering co-fluctuation patterns of varying amplitudes and examining their cluster structure at multiple hierarchical levels.

To address this question, we leveraged a hierarchical and recursive extension of the popular community detection method, “modularity maximization” [19, 20] (see **Materials and methods** for a detailed description of the algorithm). Our approach is similar to other recursive applications in that we iteratively partition sub-networks until we reach some stopping criterion. Here, we stop when the detected communities have local modularities – for each community, the sum over all within-community edges less their expected weights – that are statistically indistinguishable from a null distribution (Fig. 1f). We further excluded small communities (fewer than five elements) and those comprised of co-fluctuation patterns from only one scan session.

**FIG. 1.**
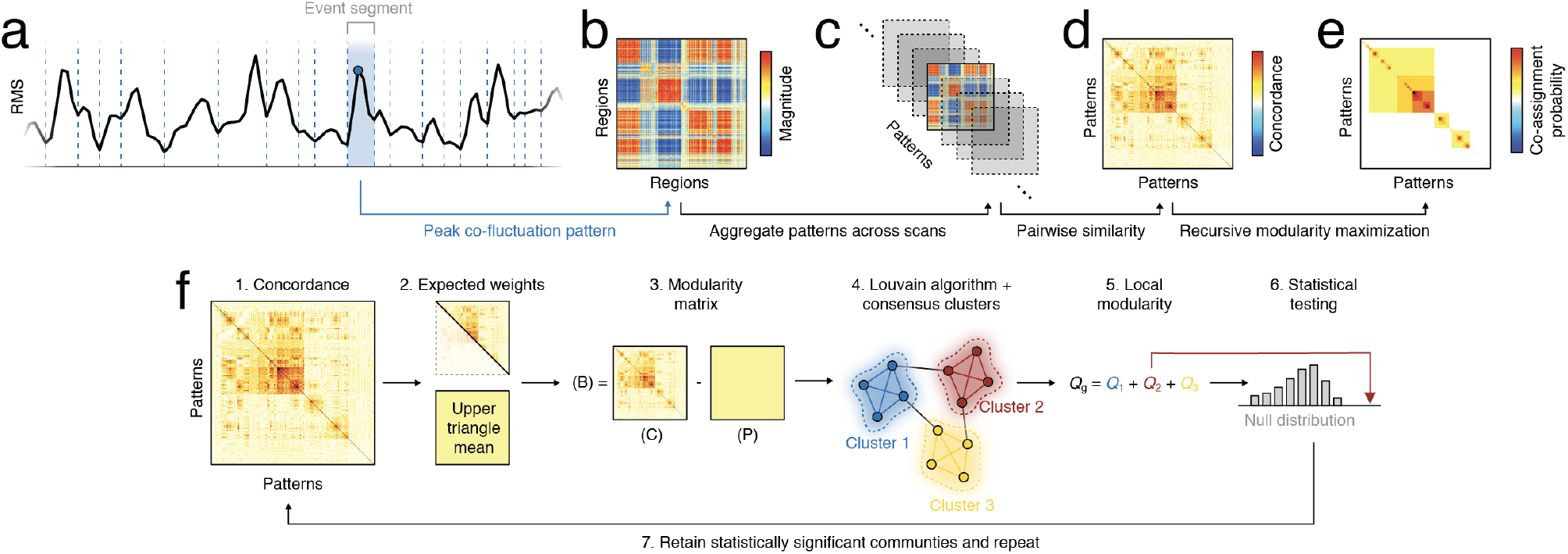
Analysis pipeline. (*a*) The global amplitude of edge time series (root mean square; RMS) was segmented into motion-free trough-to-trough intervals, or “event segments”. (*b*) Each segment was represented by the co-fluctuation pattern at its peak. (*c*) These patterns were aggregated across scans. (*d*) The similarity between pairs of co-fluctuation patterns was measured with Lin’s concordance. (*e*) A hierarchical variant of modularity maximization was used to estimate consensus community structure at different scales (resolutions). The multi-scale communities were then summarized using a matrix of co-assignment probabilities. (*f*) Detailed schematic of recursive clustering algorithm. From the full concordance matrix we estimate an expected weight as the mean of this matrices’ upper triangle elements. We calculate a modularity matrix as the observed concordance minus expected weight, and submit this to the Louvain algorithm, which we run 1000 times (different initial conditions) whose outputs are delivered to a consensus clustering algorithm. We calculate the modularity contribution (local modularity) of each consensus cluster and compare those values to a null distribution generated by randomly permuting consensus community assignments. We retain only those communities whose local modularity is statistically greater than that of the null distribution. The concordance matrices for each such community is returned to step 1 and the algorithm is repeated. This process terminates when the local modularities of all detected consensus communities are consistent with their respective null distribution.

Here, we apply this algorithm to the similarity matrix estimated from 1568 peak co-fluctuation patterns. Note, that as a measure of similarity we used Lin’s concordance in place of the more common Pearson correlation [21]. This concordance measure has been applied previously to brain network data [22] and, in contrast with Pearson’s correlation, which assesses the similarity of two patterns irrespective of their amplitudes, the concordance measure penalizes the similarity score if the amplitudes are mismatched (see **Materials and Methods** for more details).

We found that the hierarchical clustering algorithm detected ten distinct hierarchical levels (Fig. 2 *a*). At the coarsest scale (hierarchical level 2), we identified three large communities (Fig. 2a), each formed by cohesive patterns of co-fluctuations (Fig. 2b-d). The largest of these communities (cluster 1) contained 835 patterns (53.3% of all patterns), while the next largest - clusters 2 and 3 - contained 231 and 200 patterns (14.7% and 12.8% of all patterns).

**FIG. 2.**
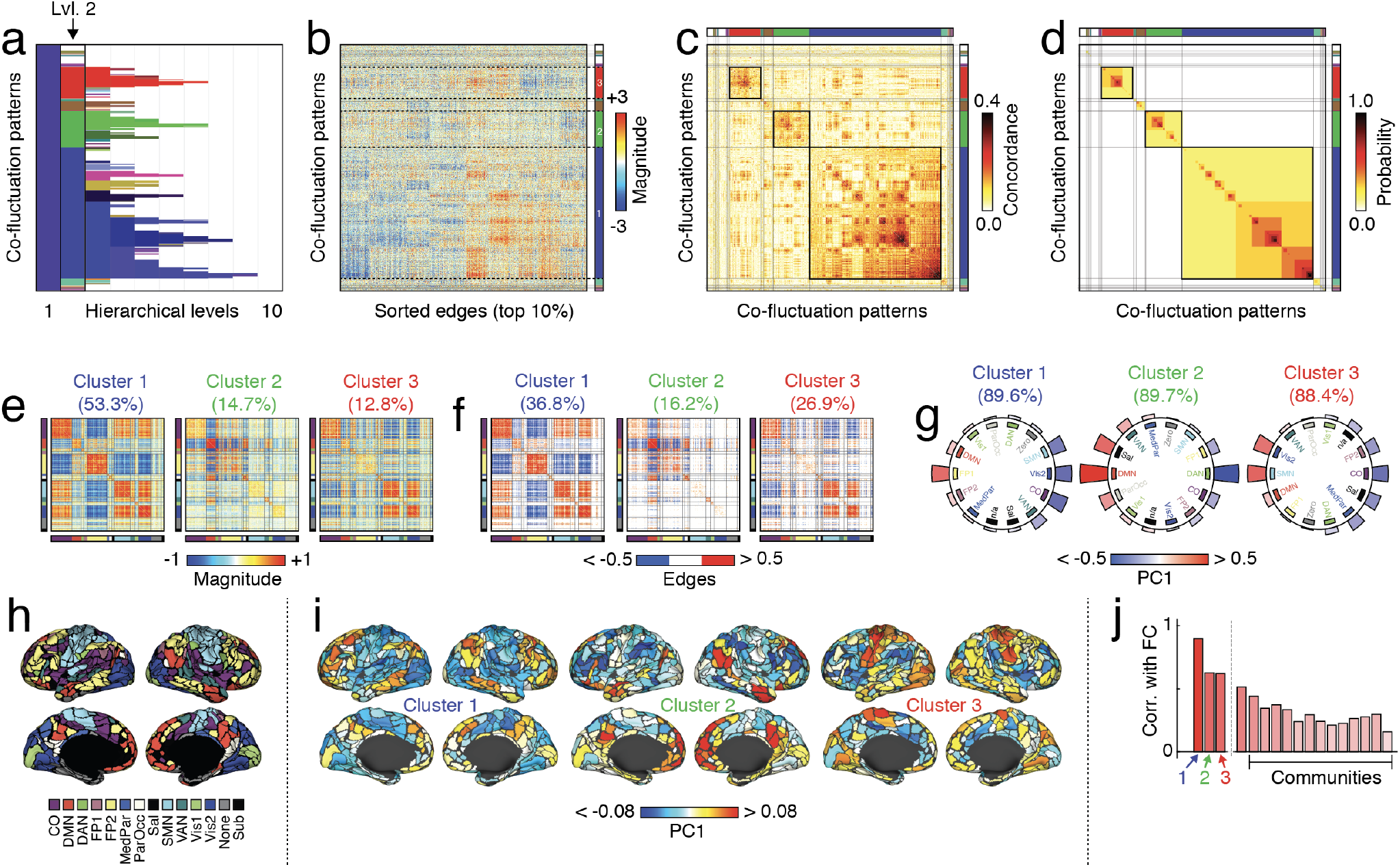
Hierarchical organization of co-fluctuation patterns. Peak co-fluctuation patterns were clustered using a hierarchical analog of modularity maximization. (*a*) Cluster labels at each hierarchical level. The color associated with each cluster was determined by first assigning each level-3 cluster a unique color (an RGB triplet) using the MATLAB function `distinguishable_colors` (https://www.mathworks.com/matlabcentral/fileexchange/29702-generate-maximally-perceptually-distinct-colors). For clusters at all other hierarchical levels, we projected their centroids onto the level 3 cluster centroids, rescaling each projection magnitudes so that, collectively, they summed to unity. Finally, we assigned each cluster a color as the linear combination of level-3 RGB triplets, each weighted by the corresponding normalized projection magnitude. (*b*) Top 10% edges, (*c*) concordance matrix, and (*d*) co-assignment matrix sorted by community label. (*e*) Cluster centroids for largest clusters at second hierarchical level. (*f*) Strongest edges in centroid matrices. (*g*) We calculated the mean value of PC1 for each brain system and plotted these values on the perimeter of a circle. In each plot the order of systems differs; those with positive co-fluctuation appear on the left and those with negative on the right. (*h*) MyConnectome system labels from [15]. (*i*) To identify dominant mode of activity underlying each cluster we calculated the first principal component (PC1) of each cluster centroid matrix. Here, we show PC1 projected onto the cortical surface. (*j*) Correlation of mean co-fluctuation patterns with static FC.

Next, we characterized these three clusters in greater detail. For each cluster, we computed its centroid as the mean over all patterns assigned to that cluster (Fig. 2e,f). In previous work, we showed that at a timescale of individual frames, co-fluctuation patterns estimated from edge time series can always be viewed as a bipartition of the network into two groups that correspond to collections of nodes whose instantaneous activity levels are above or below their respective means [5]. Even at this coarse scale, where clusters represent the average of many individual co-fluctuation patterns, the underlying bipartitions were still apparent. For example, consider cluster 1 (the largest of the three clusters). It is typified by opposed co-fluctuations of cingulo-opercular, visual, attention, and somatomotor networks (“group A”) with default mode and frontoparietal networks (“group B”) (Fig. 2e). That is, were we to examine regional BOLD data at points in time when this cluster is expressed, we would expect to find the activity of regions in groups A and B to have opposite sign (Fig. 2i). Interestingly, cluster 1, which appears most frequently, also has the strongest correspondence with static (time averaged) FC (Pearson correlation of *r* = 0.90; Fig. 2j), suggesting that the prevalence of this activity mode (and corresponding co-fluctuation pattern) help to “lock in” the gross connectional features of FC.

Clusters 2 and 3 corresponded to opposed cofluctuations of default-mode with fronto-parietal networks (cluster 2) and sensorimotor systems (somatomotor + visual networks) with salience and cingulo-opercular networks (cluster 3). These two communities were also related to static FC, albeit not as strongly correlated (*r* = 0.63 and *r* = 0.62, respectively). Note that the remaining communities, including much smaller communities, were correlated with FC but to a much lesser extent (*r* = 0.30 ± 0.09).

We also performed a series of supplementary analyses. First, to ensure that differences in the correspondence between cluster centroids and static FC was not driven by differences in the amount of data used to estimate each centroid, we repeated this analysis using individual co-fluctuation patterns. In general, the results of this analysis were consistent with those reported here; patterns assigned to cluster 1 were more strongly correlated with FC compared with those assigned to different clusters (Fig. S6). Additionally, we repeated this entire enterprise using MSC data and, despite different acquisition parameters and amounts of data, again observed consistent results (see Fig. S7, Fig. S8, Fig. S9).

As a final supplementary analysis, we assessed to what extent these results depend on our decision to cluster peaks of the RMS signal as opposed to nearby, but off-peak, frames. To address this question, we resampled the full set of 1568 peak co-fluctuation patterns by selecting frames from within the same trough-to-trough segment but with a random offset (Fig. S3a). Using these off-peak frames, we calculated a null concordance matrix that we compared with the observed matrix. We repeated this process 1000 times and, for every pair of co-fluctuation patterns, estimated the probability that their null concordance was at least as large as the observed. In general, we found that stronger concordance values corresponded to small p-values (Spearman’s rank correlation; *ρ* = −0.87, *p* < 10^-15^; Fig. S3e). That is, if two peak co-fluctuation patterns were highly concordant, any movement away from their respective peaks resulted in decreased concordance. Because high-concordance pairs tended to be assigned to the same cluster, small p-values were concentrated within clusters Fig. S3d), suggesting that the detected clusters would be systematically disrupted had we elected to cluster non-peak frames.

Collectively, this analysis of the coarsest level of cofluctuation patterns generates clusters whose centroids are consistent with those reported in our previous paper [7]. However, unlike that study, the hierarchical clustering algorithm used here allows us to investigate increasingly refined and more exclusive communities. We explore these communities in the next section.

### Sub-divisions of coarse-scale community structure

Our hierarchical clustering approach generates a nested and multi-scale description of peak co-fluctuation patterns. In the previous section we focused a single hierarchical level (the coarsest non-trivial partition). Here, we investigate the “children” of those coarse “parent” clusters. For practical reasons, we focus our investigation on sub-divisions of clusters 1 and 2 from hierarchical level 2 (described in the previous section). Comparing clusters across hierarchical levels necessitates a naming convention that not only distinguishes a cluster from other clusters in its own hierarchical level, but indicates the level in which it was detected. We now refer to clusters using the convention `hierarchical_level.cluster_number`. So cluster 1 in hierarchical level 2 would be referred to as “cluster 2.1”. Note that cluster numbers are reused across levels, i.e. cluster label 1 will appear in all layers, but will be distinguishable by the prefixes 2.1, 3.1, 4.1, and so on.

We find that cluster 2.1, which we described in the previous section, fragments into five distinct communities in the third hierarchical level (Fig. 3a-c). The first and largest of these communities, cluster 3.1, represents a refined version of its parent (Fig. 3d) in which positive and negative co-fluctuations are reinforced, strengthening their weights. In fact, of the five sub-clusters, this one maintains the strongest similarity to its parent (*r* = 0.99). Incidentally, we observe similar behavior for all three of the large clusters detected in hierarchical level 2. That is, we find evidence of sub-clusters that strongly resemble their parent, but simply increase the magnitude of the strongest positive and negative co-fluctuations (see Fig. S10).

**FIG. 3.**
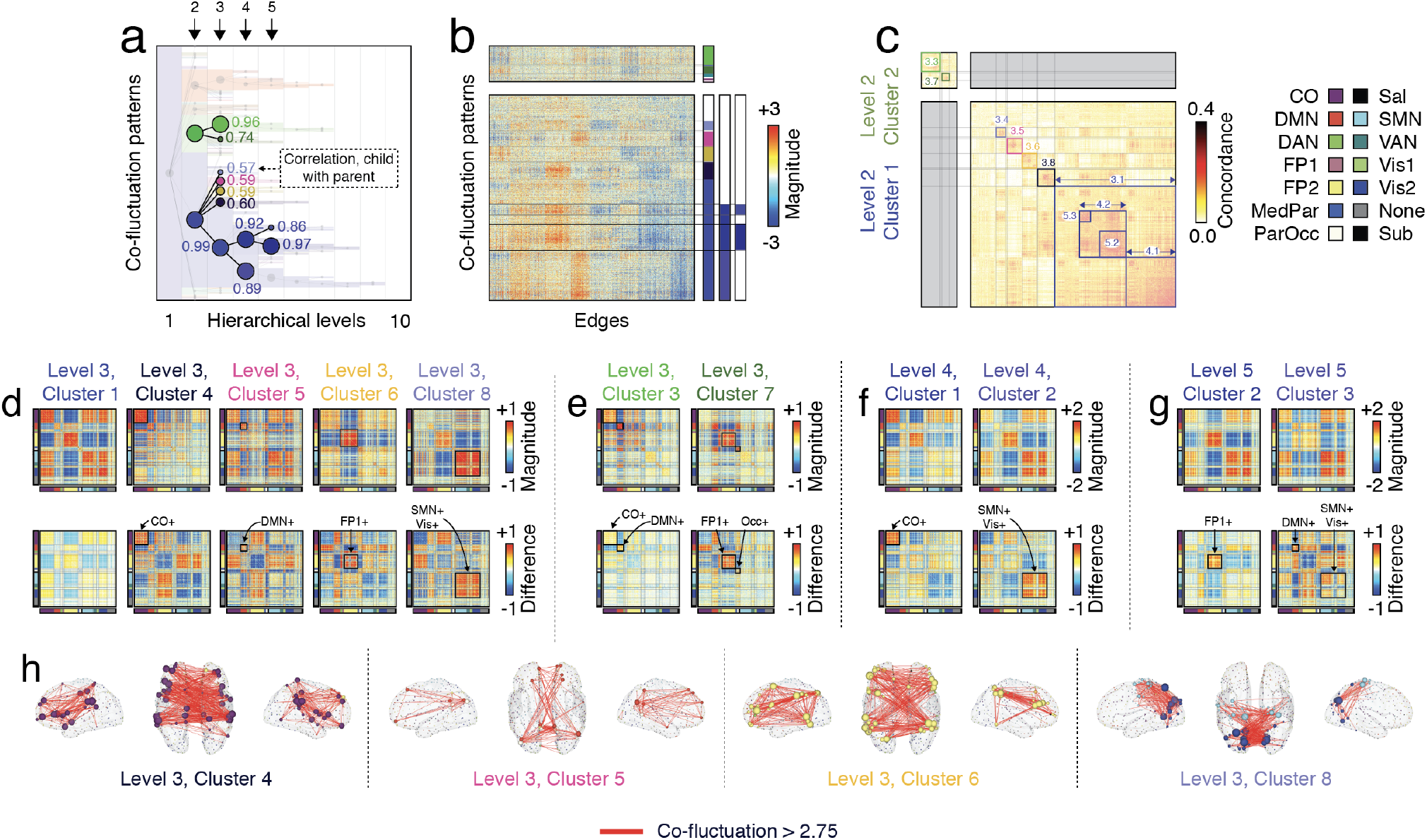
Exploring hierarchical relationships among communities. Previously we investigated a specific hierarchical level of community structure (*a*). Here, we investigate sub-divisions of those communities, focusing on what had previously been termed “cluster 1” and “cluster 2” at the second hierarchical level (we refer to these as clusters 2.1 and 2.2 following the convention `hierarchical_level.cluster_number`). We overlap a dendrogram over the community label matrix to highlight the divisions of those communities at levels 3, 4, and 5. The text next to each node in the dendrogram denotes the correlation of the corresponding co-fluctuation pattern with its parent. (*b*) A zoomed-in version of edge co-fluctuation weights for clusters 2.1 and 2.2 at the second hierarchical level, highlighting their subdivisions. (*c*) Concordance matrix ordered by communities. (*d*) Mean co-fluctuation patterns (centroids) for subdivisions of cluster 2.1 at level 3. The top row depicts sub-cluster centroid and the bottom row depicts the mean difference of children co-fluctuation patterns with their respective parents. Panels *e-g* depict analogous matrices for sub-divisions of clusters 2.2, 3.1, and 4.1. (*h*) Edges with the strongest co-fluctuation magnitude (> 2.75) for clusters 3.4, 3.5, 3.6, and 3.8.

The next four clusters (3.4, 3.5, 3.6, and 3.8), however, reflect distinct sub-components of their mutual parent. Specifically, each cluster exhibits strengthened co-fluctuations within specific functional systems. For instance, cluster 3.4 corresponds to strengthened co-fluctuations among cingulo-opercular regions, whiles clusters 3.6, 3.7, and 3.8 correspond to increases among default mode, frontoparietal, and the sensorimotor complex (comprised of visual and somatomotor systems). Notably, these sub-divisions maintain a weaker correspondence with their parent (mean correlation of *r* = 0.59 ± 0.01). We show the top co-fluctuations (edges) for these four clusters in Fig. 3h.

Interestingly, cluster 3.1 underwent further refinement in hierarchical levels 4 and 5. Clusters 4.1 and 4.2 reflect increased coupling of cingulo-opercular and a somatosensory complex to themselves, respectively (Fig. 3f), while clusters 5.2 and 5.3 split cluster 4.2 into distinct communities that reflect increased coupling of fronto-parietal regions to themselves and, separately, the default mode and sensorimotor complex to themselves (Fig. 3g).

We also found that cluster 2.2 could be further subdivided. At the coarsest level, this cluster corresponded to opposed co-fluctuations of default mode regions with cingulo-opercular and dorsal attention regions. Of its two sub-clusters, the first (cluster 3.3) could be considered a refinement and continuation of the previous coarser cluster. This cluster also maintained a strong correspondence with its parent (*r* = 0.96). In contrast, cluster 3.7 decoupled the fronto-parietal network from the default mode and cingulo-opercular systems (Fig. 3f).

The hierarchical clustering framework also allowed us to explore the composition of clusters at different levels. Specifically, we assessed the extent to which clusters were composed of high-, middle-, or low-amplitude co-fluctuation patterns. We found that, at finer scales, high-amplitude “events” acquired a greater share of the detected clusters (Fig. 4a). This observed effect belied a more general and continuous relationship between the similarity of co-fluctuation patterns to one another and their RMS. We found that high-amplitude co-fluctuation patterns tended to have greater levels of similarity to other high-amplitude frames compared to lower-amplitude frames (ANOVA, *F*(2) = 108.8, *p* < 10^-15^; post-hoc *t*-tests comparing high-amplitude frames to middle- and low-amplitude frames, *t*(1527) = 13.2 and *t*(1311) = 7.1, maximum *p* = 2.3 × 10^-12^; Fig. 4b).

**FIG. 4.**
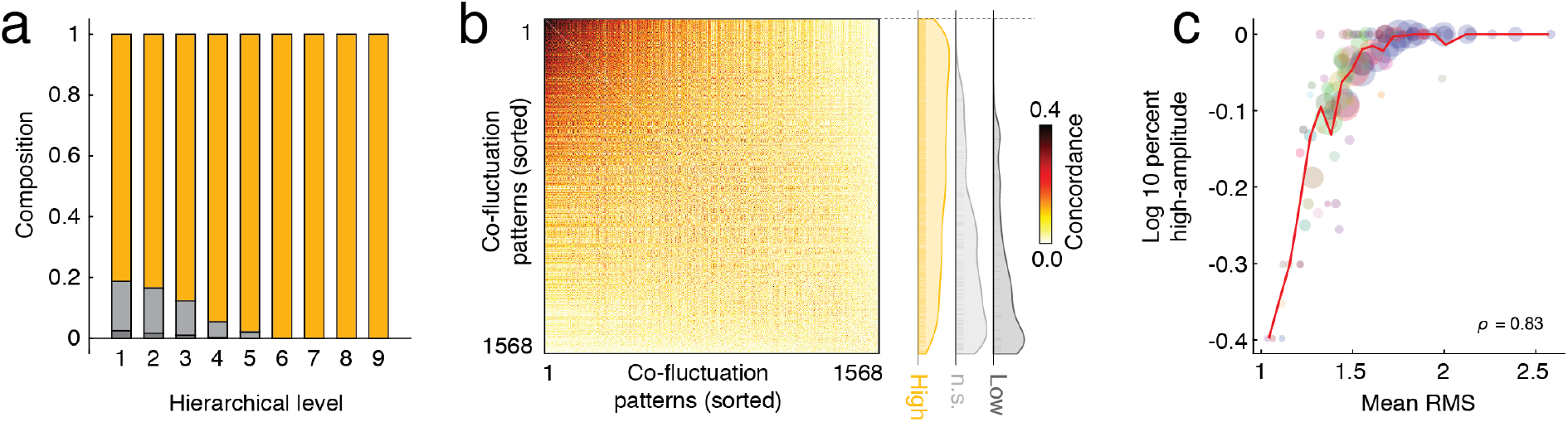
Cluster composition and similarity. At each hierarchical level, we identified the “peak type” of all frames that were not pruned away by the hierarchical clustering algorithm (“high-amplitude”, “not significant” (n.s.), or “low-amplitude”). (*a*) Composition of all patterns by peak type. Note that as clusters become more exclusive they tend to be dominated by high-amplitude frames. (*b*) Concordance matrix ordered by mean similarity of co-fluctuation patterns to one another (greatest to least). The panel to the right displays the relative positions of high-/low-amplitude and not significant patterns. Note that the low-amplitude and not significant patterns are concentrated near the bottom. (*c*) Assigning patterns a peak type is a discretization of patterns’ RMS values. Here, we show that fraction of frames assigned to a given cluster labeled high-amplitude is tightly correlated with the cluster’s mean RMS. The size of points is proportional to the number of patterns assigned to that community.

Note that the infrequency of low-amplitude peaks is due, in part, to the fact that low-amplitude frames were more likely to be censored due to high levels of in-scanner movement and thresholding based on relative RMS, but also due to a selection bias (we only sampled *peak* cofluctuations, which are necessary local maxima with RMS greater than most non-peak frames; see Fig. S4 for similarity of peaks to temporally proximal frames).

We note that we also repeated this analysis using network “templates” described in [5]. See Fig. S11.

Collectively, these observations expose the rich, multiscale and hierarchical organization of peak co-fluctuation patterns. These findings elaborate upon the clusters disclosed in the previous section and earlier papers.

### Module statistics

One of the primary aims of this manuscript was to investigate clusters of peak co-fluctuations to better understand, specifically, the contribution of low- and middle-amplitude peaks. To address these questions, we analyzed MyConnectome data - a dense sampling study of a single brain [15]. One of the advantages of analyzing so much data from a single individual is that we can assess how much data - in terms of time and scan sessions - is necessary to accurately estimate network properties. Previous studies have used these same data to understand how data is required to estimate static FC. Here, we take an analogous approach so that we can better understand how much data is required to estimate cluster centroids.

To do this, we iteratively split the complete dataset (84 scans) into two random subsets comprising 42 scans each. We then select one scan at random from one of the subsets, and, using only those data, estimate centroids for each of the clusters detected using the full set of data. We then compare those centroids to those estimated using all of the data in the other subset. We then repeat this process after we add in a second scan’s worth of data, then a third, a fourth, etc., until we have incorporated all of the data available in both subsets. This entire process then gets repeated using a different random bipartition of scans.

This procedure allows us to estimate how much data is required to achieve a fixed similarity value. First, we measure the amount of data in units of scans. With the exception of the largest cluster, the similarity curves for the smaller clusters never clearly asymptote (Fig. 5a). That is, even after 40 scans, we would expect there to be non-trivial levels of variability in our estimates of cluster centroids.

**FIG. 5.**
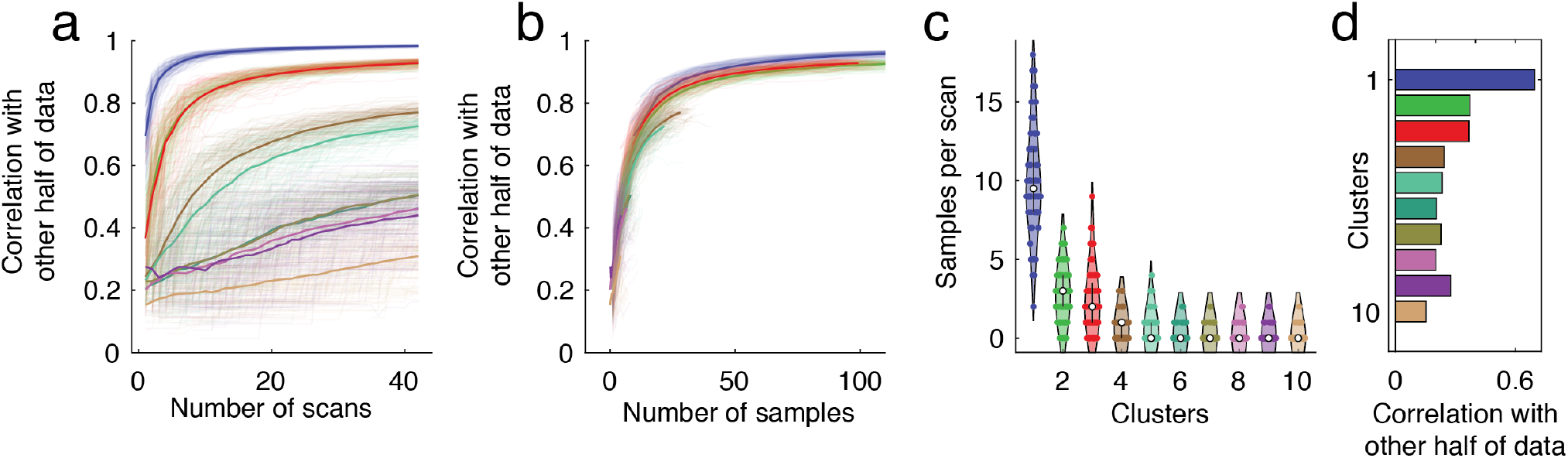
Amount of data required for accurate estimates of cluster centroids. We repeatedly and randomly split the 84 scans into two groups (42 scans each). For group 1 we used all available data to estimate cluster centroids. For group 2, we estimated centroids using data from one random scan and sequentially incorporated data from additional scans. At each step, we calculated the similarity of cluster centroid estimates from group 2 with estimates from group 1. (*a*) Similarity for clusters 1-10 and hierarchical level 3 as a function of number of scans. (*b*) Similarity as a function of samples. (*c*) Number of times that each cluster appears in a given scan. (*d*) To further control for differences in the number of samples and to assess whether some clusters were composed of patterns that were inherently more similar to their mean, we calculated the mean correlation of individual co-fluctuation patterns in one half of the data with the centroids from the other half.

To understand why this happens, we need to change our unit for measuring the amount of data from “scans” to “samples” and also estimate the baseline frequencies with which each cluster type occurs. In general, we find that co-fluctuation patterns labeled as cluster 1 occur, on average, 9.9 ± 3.4 times per scan (Fig. 5c). The next most frequently occurring cluster appears only 2.75 ± 1.6 times per scan. In other words, we expect that each additional scan would yield ≈10 new instances of cluster 1 but between 2 and 3 instances of cluster 2. Therefore, if our aim were to acquire a fixed number of samples of a cluster 2 compared to cluster 1, we would require proportionally four times as many scans. Indeed, when we recreate Fig. 5a where units are now in number of samples rather than number of scans, we find that the similarity curves for all clusters overlap (Fig. 2b).

Collectively, these results suggest that a key limiting factor in accurately estimating cluster centroids is the relatively low frequencies with which some of the smaller states occur. Our results also suggest that a key factor contributing to variability in connectivity patterns from one day to another might be the frequency with which different cluster patterns appear.

### Linking FC and hierarchical depth

The hierarchical procedure yields a progressively sparse perspective on recurring co-fluctuation patterns, with higher levels of the hierarchy appearing more exclusive and containing progressively fewer co-fluctuation patterns but ones that form extremely tight and cohesive clusters (Fig. 6a). Because previous studies have linked co-fluctuation patterns to FC, this hierarchical perspective allows us to assess at what hierarchical level (and by extension, what level of exclusivity) do co-fluctuation patterns most closely correspond to FC.

**FIG. 6.**
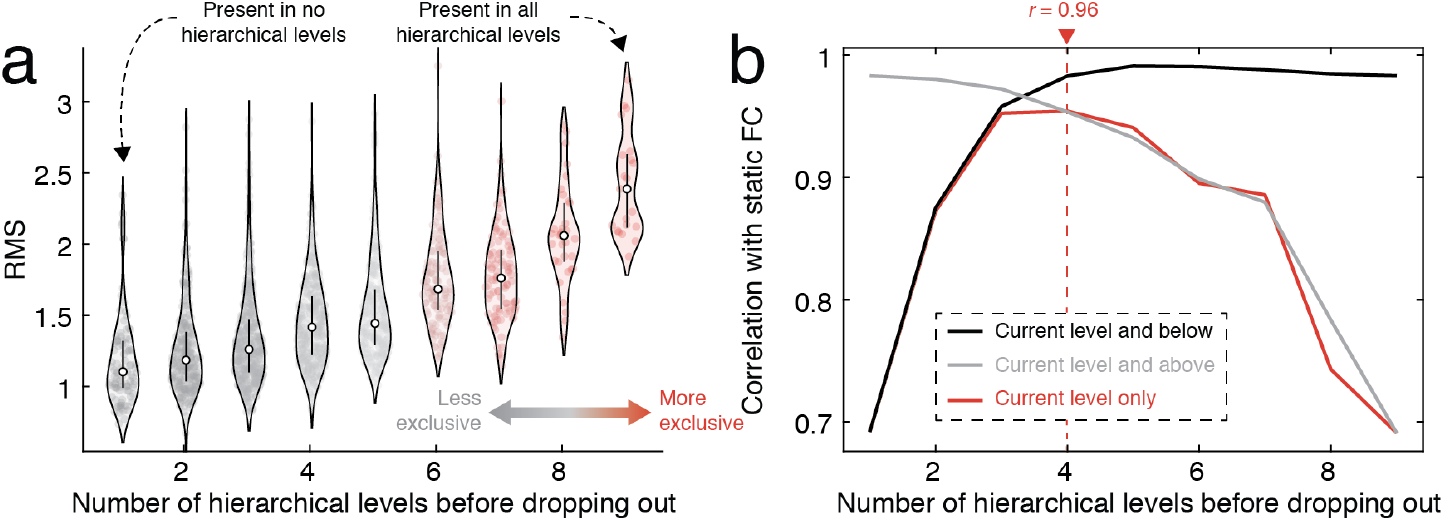
Linking hierarchical levels to FC. We calculated the deepest level of the hierarchy to which each co-fluctuation pattern was assigned (the number of levels in which it appeared before being pruned away). (*a*) RMS of co-fluctuation patterns, grouped by hierarchical depth. (*b*) The correlation of mean co-fluctuation patterns with FC. We compared three estimates of the mean: using the current level and below, the current level and above, and the current level only.

To address this question, we assigned each cofluctuation pattern a score indicating its “hierarchical depth.” That is, the number of hierarchical levels in which that pattern was present and included in a community. Then we followed the procedure outlined in [7] where, separately for each hierarchical level, we calculated the average co-fluctuation pattern across all patterns assigned to that level, and computed the correlation of that matrix with static FC (based on their upper triangle elements). We also repeated this procedure using a cumulative approach (starting with patterns at the highest level, gradually incorporating patterns from lower levels until all patterns were included) and a reverse cumulative approach (starting with the coarsest scale, gradually peeling away coarser and coarser partitions until only the most exclusive patterns were included).

In general, we found a statistically significant correspondence between FC and co-fluctuation patterns at all hierarchical levels using all three methods for estimating the mean co-fluctuation pattern (minimum *r* = 0.69; *p* < 10^-15^). Using patterns from individual levels only, we found that this correspondence peaked at the fourth hierarchical level (*r* = 0.96), suggesting that the correspondence with FC is maximized when including not only the highest-amplitude co-fluctuations, but weaker and less exclusive communities as well. We observed a similar trend using the cumulative and reverse-cumulative approaches – starting with exclusive clusters and including patterns assigned to less-exclusive clusters led to improvements in the correspondence, as did starting with all patterns and pruning away weaker patterns. We find similar results with MSC data (see Figure. S12).

Collectively, these observations build on results from our previous studies, noting that high-amplitude cofluctuations are indeed correlated with FC, but that this correlation can be improved upon by including more heterogeneous and slightly weaker co-fluctuation patterns. That is, the lower-amplitude patterns, which are more variable, effectively “sculpt” the more stereotypical cofluctuation pattern driven by high-amplitude events, enhancing the diversity of co-fluctuations and improving the correspondence with static FC. Moreover, we find that the correspondence is maximized at an intermediate level, suggesting that different hierarchical scales are differentially informative about FC.

## DISCUSSION

In this paper we build on previous studies of edge time series. Namely, we focused on the statistics of peaks in the RMS time series and the clusters formed by the co-fluctuation patterns expressed during the peaks. We developed a bespoke multi-scale clustering algorithm to construct a hierarchy from peak co-fluctuation patterns and investigated clusters at all scales, ranging from coarse clusters that included most patterns to exclusive clusters composed of only a small number of patterns. Finally, we assessed how the amount of data impacts estimates of clusters and their centroids. Collectively, this work addresses several gaps in knowledge and further demonstrates the utility of edge analyses for fMRI data.

### Co-fluctuation patterns are hierarchically organized in time

A large body of work has shown that nervous systems exhibit multi-scale, hierarchical organization [23, 24]. Overwhelmingly, this work has focused on hierarchical *spatial* structure, in which neural elements are organized into modules within modules within modules, *ad infinitum* [25–29].

In contrast, there are fewer papers that focus on hierarchies in time. This is not to say that the temporal organization of nervous systems – and functional brain networks, in particular – has gone uncharacterized. In fact, the opposite is true; time-varying connectivity analyses have come to occupy an increasingly large share of contemporary network neuroscience and connectomics [30-36] (despite several papers that cast doubt on the very premise that network change can be measured with fMRI [37, 38]).

One of the key findings in the time-varying connectivity literature is that brain networks appear to traverse a series of “network states”, i.e. a pattern of connectivity approximately persists for some period of time before giving way to a new pattern of connectivity. There also exists mounting evidence that these states are revisited across time within an individual and shared across subjects at the population level [39–43].

In most state-based analyses, time-varying estimates of network structure are usually obtained from sliding-window methods. The sliding window approach, however, represents only one strategy for obtaining (smoothed and temporally imprecise) estimates of timevarying connectivity ([44–46]. Recently, we proposed a method for tracking “instantaneous connectivity” across time, obviating the need for sliding windows [3, 4] ^1^. In contrast with the smooth variation observed using sliding windows, edge time series exhibited “bursty” behavior – long periods of quiescence punctuated by brief high-amplitude events. The co-fluctuation patterns coincident with events were strongly correlated with static FC (aligned with earlier findings [10, 12-14]), contained subject-identifying features, and, in an exploratory analysis, strengthened brain-behavior correlations [3]. Note that in these first studies, we treated edge time series as an instantaneous estimate of time-varying connectivity. However, edge time series can be viewed more generally as a decomposition of any the correlation between any two variates, irrespective of whether they are, in fact, time series.

Following our initial work, we showed that high-amplitude co-fluctuations could be partitioned into at least two distinct clusters or “states.” These states were shared at the group level, but refined individually, leading to the personalization of subjects’ FC patterns [6, 7]. This two-state description, however, was a direct consequence of the method used to detect states and precluded the possibility that co-fluctuation patterns were organized into clusters at multiple scales or hierarchically. This type of temporal organization, in which broad “meta-states” could be sub-divided into a series of smaller states has been observed in other contexts, e.g. clusters of independent components [48–50] or states estimated using hidden Markov models [51]. However, it was unclear whether edge time series exhibited analogous temporal structure.

To address this question, we investigated data from the MyConnectome project. Our rationale for selecting this dataset was that, with > 10 hours of resting-state data, the MyConnectome project gives us the best chance to detect infrequent states, if they exist, and to obtain better estimates of the states that occur more frequently. to investigate co-fluctuation states, we developed a hierarchical clustering algorithm built upon recursive application of the familiar modularity maximization algorithm [19], allowing us to obtain estimates of large “metastates” but also smaller, more refined states.

We found evidence of three large clusters of cofluctuation patterns that persisted over multiple hierarchical levels, gradually refining their organization. Notably, the centroids of these clusters were consistent with those reported in our previous work [7], and were aligned with other recent findings. For instance, our clusters delineate task-positive and -negative systems [52, 53], recapitulate spatial modes of variation in resting-state data time series [54], and closely resemble components of so-called “functional gradients” [55], which are frequently interpreted in terms of cognitive hierarchies [56].

Importantly, we also found that these large clusters could be meaningfully sub-divided into smaller, increasingly nuanced patterns of co-fluctuation. This observation has important implications for how we interpret static FC, but also for our understanding of brain dynamics and inter-areal communication. Because edge time series are an exact decomposition of FC into framewise contributions, the average across peak co-fluctuations serves as an approximation of FC. While this pattern-level estimate is generally very accurate, it offers no compression (each individual pattern is needed). By clustering patterns we reduce the description length of co-fluctuation patterns while, hopefully, still generating a good approximation of FC. Indeed, we find that this is the case, with different hierarchical levels and clusters offering variable predictions. Of particular interest is the observation that mid-hierarchy co-fluctuations actually outperform other levels. This is because clusters of co-fluctuation patterns at coarse scales are too general to recapitulate details of FC connectivity patterns, while clusters at the finest scales are too specific.

FC is frequently interpreted as evidence of communication or coordination between pairs of brain regions [2, 57]. This interpretation is evident when we consider the brain’s static system organization - i.e. its division into subnetworks like the default mode, visual, and attentional systems - we generally think of these cohesive modules as reflecting the outcome of a segregated and functionally specialized process. In previous studies, however, we demonstrated that, as measured with fMRI, no more than two modules can be “engaged” at any single point in time, implying that the brain’s static system-level architecture is a consequence of dynamically fluctuating bipartitions that, occasionally, do not resemble any of the frequently-discussed brain systems [5]. Moreover, if we think of static FC as a reflection of inter-areal communication, then each bipartition - and especially those that occur during peaks, when the co-fluctuation magnitude is much stronger than nearby frames - may reflect a communication event. Our findings, here, suggest that these instants of communication are highly structured in space and time. Spatially, we identify a richer repertoire of cofluctuation patterns than had previously been reported (more clusters) and show that, while these patterns can involve the entire cerebral cortex, they also can engage specific subsets of systems. Our findings also suggest that these patterns occur intermittently but recur across time. Thus, the brain’s temporal trajectory as defined by edge time series is low-dimensional, but also bursty.

### Hierarchies contain heterogeneous co-fluctuation patterns of similar amplitude

In most previous analyses of edge time series, emphasis was placed on high-amplitude frames [3, 6, 7]. That is, instants in time where the global co-fluctuation amplitude was disproportionately large. The rationale for doing so was that, because FC is literally the mean of an edge time series, frames with large amplitude *must* contribute more to the average and frames where many edges have large amplitude necessarily contribute more to the overall FC pattern. However, these high-amplitude periods are rare and while, on a per frame basis, they contribute more than middle and lower amplitude frames, they number far fewer. Moreover, they represent only the tail of a distribution and ignore low-amplitude frames, which tend to be more susceptible to motion artifacts [7], but also middle-amplitude frames, about which less is known.

Here, we find that the overall magnitude of cofluctuations scales with hierarchical level. That is, the co-fluctuation patterns that make up the most exclusive and highest level of the hierarchy tend to be composed of those with the greatest overall amplitude, while lower-amplitude patterns populate the intermediate levels of the hierarchy. This observation is analogous to recent findings, reporting a graded link to FC [58]. However, our findings also suggest that nuance is necessary in describing links between amplitude and FC and that, at every hierarchical level, there exists structured heterogeneity of co-fluctuation patterns, i.e. they can be grouped into clusters.

### Accurate estimates of cluster centroids require lots of data

One of our key observations is that, if we want to accurately estimate cluster centroids, we require large amounts of data. For some of the smaller and less frequently appearing clusters, this amount is prohibitively large and infeasible for most fMRI studies (greater than 7 hours). This observation is in line with other studies showing that a major source in the variability of functional brain networks is the amount of data [15, 18, 59]. In fact this effect gets amplified when estimating cofluctuations; while a typical scan session samples brain activity at hundreds of time points, a much smaller fraction of those will correspond to peaks.

While this effect can be viewed as a limitation, it also serves as a potential explanation for observed variability in network architecture from one day to the next. Because FC is the average of co-fluctuation patterns across time, differences in cluster frequencies across scan sessions will, necessarily, correspond to differences in FC weights.

### Limitations and future directions

One of the limitations of this study is its reliance on “dense-sampling” datasets. The rationale for studying these types of data (rather than cross-sectional datasets) comes from our previous studies [7], where we demonstrated that recurring co-fluctuation patterns, while similar across individuals, are also individualized. Accordingly, we aimed to study co-fluctuation patterns at the individual level rather than at the cohort, where cluster centroids, because they are composed of patterns from many individuals, may not be representative of any of those individuals. However, while dense-sampling studies allow researchers to characterize individuals in great detail, they make it challenging to generalize to the population/cohort level. Nonetheless, there is value in examining effects at that level, as many populations are not amenable to dense-sampling designs, necessitating cross-sectional analysis. Future studies should extend this work to larger cross-sectional datasets, e.g. the Human Connectome Project.

Recent papers have shown that some of the apparently “dynamic” features of edge time series, including the emergence of events, can be explained parsimoniously by properties of the static FC matrix, e.g. its eigenspectrum [60, 61]. First, we note, that this does not change the view of edge time series as a decomposition of FC – the mean of an edge time series is still exactly that edge’s weight. Second, even if one were to accept that the peak co-fluctuations do not occur “dynamically” but reflect sampling variability around a stationary correlation structure, we can still view edge time series (and clusters of co-fluctuation patterns), from an explanatory perspective, analogous to how we interpret the results of a principal component analysis (where components correspond to modes of variability that explain linear dependencies in the larger dataset). While there is an indisputable mathematical equivalence between fluctuations in edge time series and static FC, there remain dynamic features that are not easily dismissed. For instance, it was observed that edge time series synchronize across individuals during movie-watching [3]; this effect is unanticipated if edge time series were stochastic fluctuations around a stationary correlation structure.

There exist other overarching philosophical disputes concerning the origins of and appropriate null models for edge time series (and task-free brain activity more generally). For instance, observed fMRI BOLD time series are generated by an underlying dynamical system constrained by anatomical connectivity [62–64]. That is, there exists an evolution operator that maps a pattern of activity at time *t* to a new pattern at time *t* + 1, and this operator is parameterized by SC (among other parameters). The activity time series generated by this dynamical system can, of course, be summarized by its correlation structure, i.e. its FC. However, FC itself plays no role in determining the evolution of brain activity in the model. That is, FC is a summary statistic, ephiphenomenal, and over short timescales plays no role in shaping the character of ongoing brain activity. Rather, brain activity is shaped by dynamics that are constrained by anatomy. However, many “null” models stochastically generate synthetic fMRI BOLD data given a fixed correlation structure, often estimated from the data itself [65], circularly presupposing that the observed correlation structure is the driver of itself. In short, while the results reported here do not directly speak to the dynamics of co-fluctuation time series, they set the stage for future studies to perform detailed explorations using generative models grounded in anatomical connectivity [66].

As part of this paper, we create or use several tools that might be useful for future studies. First, we used an existing measure of concordance [21] for assessing the similarity of co-fluctuation patterns to one another rather than correlation measures, which are far more common. Our rationale for choosing concordance is that it is sensitive to differences in amplitude. Imagine having two co-fluctuation (or connectivity) patterns – they are identical patterns but in once case, all edge weights are scaled by a number very close to zero so that, effectively, each weight is zero, but there remains a faint impression of the original co-fluctuation pattern. The correlation of these two patterns is exactly 1, despite the vast difference in amplitude. Their concordance, on the other hand, would be near 0. In short, concordance is a more conservative measure of similarity and could be applied in other contexts to assess the correspondence between connectivity or co-fluctuation matrices.

The second innovation is the multi-scale and hierarchical clustering algorithm. It addresses several limitations of community detection methods frequently applied to neuroimaging data. First, unlike single-scale community detection algorithms, it generates multi-scale estimates of communities at different resolutions. Note that there are many algorithms and approaches for generating multiscale estimates of communities including varying resolution parameters [67] or sparsity levels [68], although these approaches do not explicitly establish hierarchical relationships between scales, which our method does. Additionally, and importantly, our approach incorporates an internal null model that makes it possible to reject communities, an important consideration given that descriptive community detection methods can spuriously detect communities without proper statistical controls [69]. Here, we test the local modularity contributions of each community, retaining those where the contribution is significantly greater than that of a chance model. We note, however, that other criteria could be substituted and used to determine whether a community is propagated to the next level or not. Finally, the algorithm is computationally efficient in comparison to other similar methods [70]. Future analyses should focus on benchmarking this method.

## MATERIALS AND METHODS

### Midnight Scan Club

The description of the Midnight Scan Club dataset acquisition, pre-processing, and network modeling is described in detail in [17]. Here, we provide a high-level overview. Data were collected from ten healthy, righthanded, young adult participants (5 females; age: 24-34). Participants were recruited from the Washington University community. Informed consent was obtained from all participants. The study was approved by the Washington University School of Medicine Human Studies Committee and Institutional Review Board. This dataset was previously reported in [17, 18] and is publicly available at https://openneuro.org/datasets/ds000224/versions/00002. Imaging for each participant was performed on a Siemens TRIO 3T MRI scanner over the course of 12 sessions conducted on separate days, each beginning at midnight. In total, four T1-weighted images, four T2-weighted images, and 5 hours of resting-state BOLD fMRI were collected from each participant. For further details regarding data acquisition parameters, see [17].

High-resolution structural MRI data were averaged together, and the average T1 images were used to generate hand-edited cortical surfaces using Freesurfer [71]. The resulting surfaces were registered into fs_LR_32k surface space as described in [72]. Separately, an average native T1-to-Talaraich [73] volumetric atlas transform was calculated. That transform was applied to the fs_LR’_32k surfaces to put them into Talaraich volumetric space.

Volumetric fMRI pre-processing included slice-timing correction, frame-to-frame alignment to correct for motion, intensity normalization to mode 1000, registration to the T2 image (which was registered to the high-resolution T1 anatomical image, which in turn had been previously registered to the template space), and distortion correction [17]. Registration, atlas transformation, resampling to 3 mm isotropic resolution, and distortion correction were all combined and applied in a single transformation step [74]. Subsequent steps were all completed on the atlas transformed and resampled data.

Several connectivity-specific steps were included (see [75]): (1) demeaning and de-trending of the data, (2) nuisance regression of signals from white matter, cerebrospinal fluid, and the global signal, (3) removal of high motion frames (with framewise displacement (FD) > 0.2 mm; see [17]) and their interpolation using power-spectral matched data, and (4) bandpass filtering (0.009 Hz to 0.08 Hz). Functional data were sampled to the cortical surface and smoothed (Gaussian kernel, *σ* = 2.55 mm) with 2-D geodesic smoothing.

The following steps were also undertaken to reduce contributions from non-neuronal sources [75, 76]. First, motion-contaminated frames were flagged. Two participants (MSC03 and MSC10) had high-frequency artifacts in the motion estimates calculated in the phase encode (anterior-posterior) direction. Motion estimate time courses were filtered in this direction to retain effects occurring below 0.1 Hz. Motion contaminated volumes were then identified by frame-by-frame displacement (FD, described in [77]), calculated as the sum of absolute values of the differentials of the 3 translational motion parameters (including one filtered parameter) and 3 rotational motion parameters. Frames with FD > 0.2 mm were flagged as motion-contaminated. Across all participants, these masks censored 28%±18% (range: 6% - 67%) of the data; on average, participants retained 5929±1508 volumes (range: 2733 - 7667). Note that in this paradigm, even the worst participant retained almost two hours of data. Nonetheless, we excluded two subjects from all analyses, both of whom had fewer than 50% usable frames in at least five scan sessions (MSC08 in 7/10 and MSC9 in 5/10).

Time courses were extracted from individualized parcellations (see [78] for details). The time series were used for FC estimation and edge time series generation.

### MyConnectome dataset

All data and cortical surface files are freely available and were obtained from the *MyConnectome Project*’s data-sharing webpage (http://myconnectome.org/wp/data-sharing/). Specifically, we studied pre-processed parcel fMRI time series for scan sessions 14-104. Details of the pre-processing procedure have been described elsewhere [15, 79]. Each session consisted of 518 time points during which the average fMRI BOLD signal was measured for *N* = 630 parcels or regions of interest (ROIs). With a TR of 1.16 s, the analyzed segment of each session was approximately 10 minutes long.

### Functional connectivity

Functional connectivity (FC) measures the statistical dependence between the activity of distinct neural elements. In the modeling of macroscale brain networks with fMRI data, this usually means computing the Pearson correlation of brain regions’ activity time series. To calculate FC for regions *i* and *j*, then, we first standardize their time series and represent them as z-scores. We denote the z-scored time series of region *i* as **z**_i_ = [*z_i_*(1),…, *z_i_*(*T*)], where *T* is the number of samples. The Pearson correlation is then calculated as:

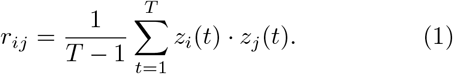

In other words, the correlation is equal to the temporal average of two regions’ cofluctuation.

### Edge time series

We analyzed edge time series data. Edge time series can be viewed as a temporal decomposition of a correlation (functional connection) into its framewise contributions. Note that Pearson correlation is calculated as 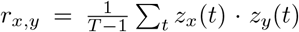, where *T* is the number of samples and 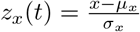 is the z-scored transformation of the time series *x* = [*x*(1),…, *x*(*T*)]. If we omit the summation in our calculation of *r_x,y_*, we obtain a time series *r_x,y_*(*t*) = *z_x_*(*t*) · *z_y_*(*t*), whose elements index the instantaneous co-fluctuation between variates *x* and *y*. Here, we estimated the edge time series for all pairs of brain regions {*i,j*}.

### Time series segmentation and peak detection

In estimating edge time series, we censored all frames with high levels of motion. In addition, we further censored frames that were within two TRs of a high-motion frame. Finally, of the remaining frames, we discarded any temporally contiguous sequences of low-motion frames that were shorter than five TRs. For a given scan session, this procedure induced discontinuous sequences of low-motion data. Aside from z-scoring parcel time series (for which the mean and standard deviation were estimated using all low-motion frames), all subsequent analyses were carried out separately for each sequence.

Let *r_ij_* (*t*) be the co-fluctuation magnitude between regions *i* and *j* at time *t*. The magnitude of cofluctuation at any instance can be calculated as 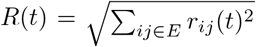 is the set of all node pairs (edges) and *N* is the total number of nodes (630 for the MyConnectome dataset). We used MATLAB’s `findpeaks` function to identify local minima in the RMS time series, resulting in 3124 low-motion, trough-to-trough segments. Within each segment there exists a single peak frame; we then calculated its relative RMS as its height minus the height largest of its neighboring troughs. We retained only those peaks whose relative RMS was greater than 0.25, reducing the number of segments to 1717. As a final exclusionary criterion, we identified peaks that occurred within 10 seconds of one another and retained only the peak with the greater relate RMS, further reducing the number of segments to 1568 (50.1% of the original).

### Lin’s concordance

In order to detect clusters among peak co-fluctuation patterns, we needed a distance metric to assess their pairwise similarity. A common candidate in human neuroimaging and network neuroscience studies is the Pearson correlation (correlation similarity). However, this measure rescales patterns before computing their similarity (z-score). That is, two co-fluctuation patterns with very different magnitude would be considered highly similar using the correlation metric. Here, however, we explicitly aimed to compare co-fluctuation patterns of differing amplitude and needed a distance/similarity metric sensitive to these differences.

Accordingly, we opted to use Lin’s concordance as a measure of similarity [21]. Briefly, this measure simultaneously assesses the similarity between two vectors based on their overall pattern (like the correlation matrix), but also allows vectors to be distinguished from another if their magnitudes differed. Briefly, the concordance between two vectors, *x* and *y*, is calculated as:

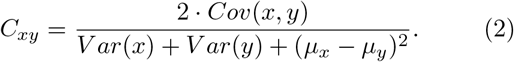

Intuitively, if the two vectors have identical means and variances, then their concordance is equal to their correlation coefficient *r_xy_*, which serves as an upper bound. However, if the variances or means of *x* and *y* differ, then *C_xy_* < *r_xy_*. Here, we calculate the pairwise concordance between all peak co-fluctuation patterns. This matrix is calculated separately for each subject.

### Recursive modularity maximization, modularity contributions, and statistical tests

We used a community detection algorithm to partition co-fluctuation patterns into hierarchically related clusters. Specifically, we recursively applied modularity maximization to the concordance matrix. The modularity heuristic defines communities as groups of nodes whose density of connections to one another maximally exceeds what would be expected by chance.

In general, the modularity, *Q,* of a partition can be expressed as the sum of contributions made by each community, *c* ∈ {1,…, *K*}, such that:

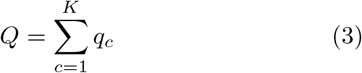

where *q_c_* =∑_*i*ϵ*c,j*ϵ*c*_[*C_ij_* – *P_ij_*]. In this expression, *i* and *j* correspond to distinct elements in the network (in our case, peak co-fluctuation patterns). The values of *C_ij_* and *P_ij_* correspond to the observed and expected concordance between those pairs of patterns.

Our algorithm is simple and built upon the modularity maximization framework. For a given concordance matrix, we uniformly set the expected weight of connections equal to the mean concordance value. That is 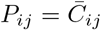 for all {*i,j*} pairs, where 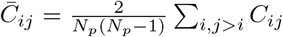 and *N_p_* is the total number of peak co-fluctuation patterns detected.

Next, we used a generalization of the Louvain algorithm [20] to optimize the modularity *Q*, repeating the procedure 1000 times with random restarts, before obtaining a consensus partition [80]. In general, the consensus partition will contain *K* communities, each of which is associated with a modularity contribution, *q_c_*. We compare the observed contribution value against a null distribution generated by preserving the consensus community labels but randomly assigning peak co-fluctuation patterns to communities (10000 repetitions). We then calculate a *p*-value as the fraction of times that the null value was greater than that of the observed value. Communities were propagated to the next level if *p* < 0. 05.

If a community survived this statistical test, it was propagated to the next level, where the entire procedure is repeated. This algorithm continues until no detected communities pass the statistical test. The end results is a series of nested communities that can be linked to one another *via* a dendrogram.

### Centroid analysis

Throughout this report we found it useful to examine individual communities in more detail. One way to summarize a community is by computing the mean cofluctuation pattern across all patterns assigned to that community, i.e. the community’s centroid. The elements of this pattern can be represented as a [*node × node*] cofluctuation matrix.

To better understand the modes of activity that underpin each co-fluctuation pattern, we performed an eigende-composition of each co-fluctuation matrix, which yielded a series of [*node* × 1] eigenvectors, each associated with an eigenvalue that was linearly proportional to the amount of variance explained by its corresponding eigenvector. We focus only on the eigenvector corresponding to the largest eigenvalue.

### Bipartition analysis

In a previous study we showed that the co-fluctuation pattern expressed at each moment in time could be partitioned into exactly two communities (a bipartition) based on whether each node’s activity was above or below its mean value [5]. These bipartitions effectively retain only the signs of a node’s activity and necessarily discard details about its amplitude. Despite this loss of information, the mean co-assignment of nodes to the same community closely recapitulate the brain’s static connectivity structure.

The bipartitions framework also facilitates a straightforward comparison with canonical brain systems. Specifically, brain systems, e.g. default mode, visual, control networks, etc., can be represented as bipartitions. The nodes assigned to that system are given a value of ‘1’ while the others are assigned ‘0’. One could also consider combinations of systems, e.g. by assigning nodes in systems A + B + C a value of ‘1’. The empirical and system bipartitions can be compared to one another directly using the measure of normalized mutual information – values close to 1 indicate a correspondence between the two bipartitions; values near 0 indicate no relationships.

Here, we used the bipartition analysis to relate each peak co-fluctuation pattern with one of 127 template patterns (all possible combinations of the 14 systems).

---

**FIG. S1.**
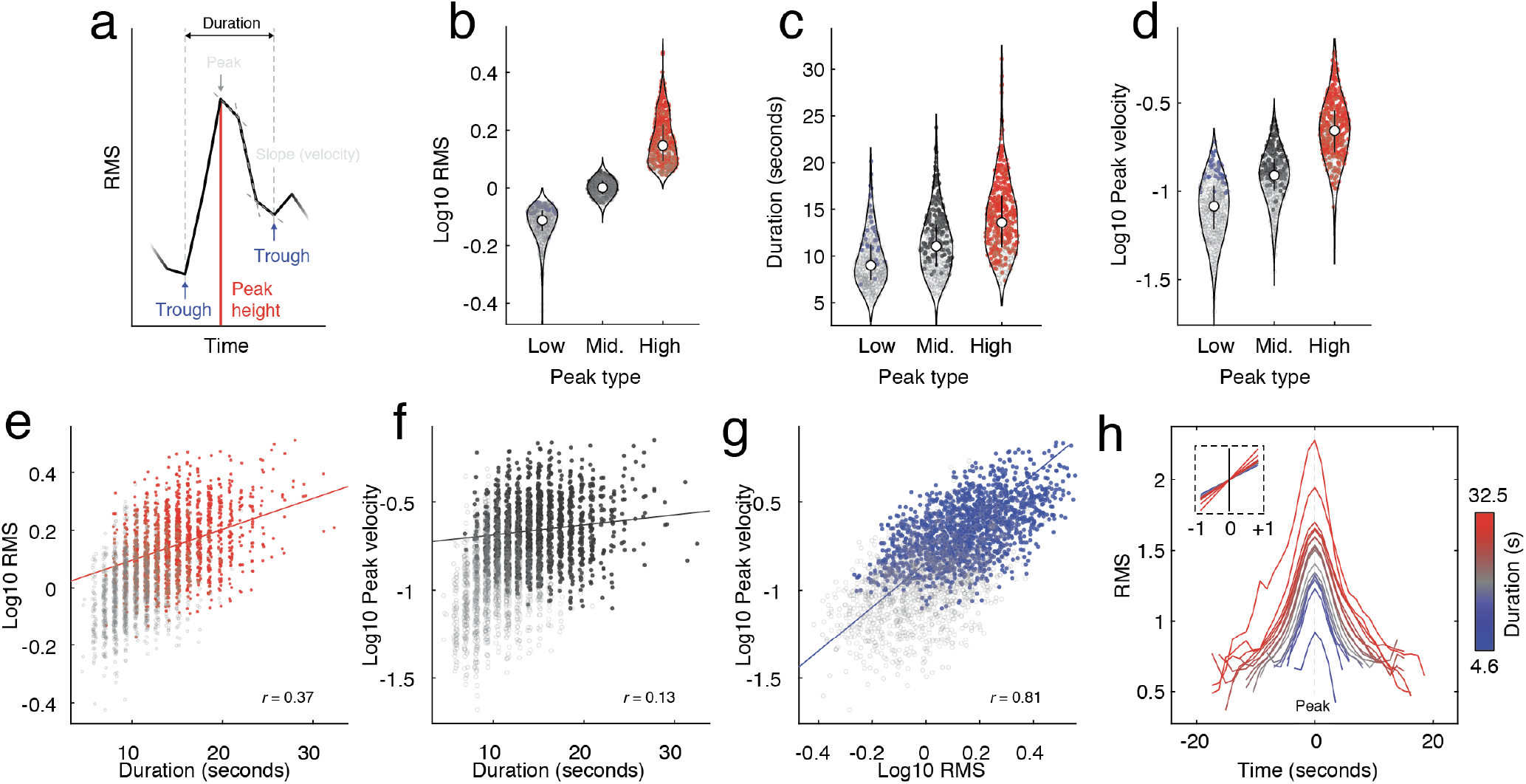
Characterizing peak co-fluctuations. For every peak, we calculated its amplitude (RMS) and duration. Using a procedure developed in our previous work, also classified each peak as a high-, middle-, or low-amplitude frame. In the main text and in all panels we report statistics based on a set of 1568 co-fluctuation patterns that survived a series of quality assessments to reduce the likelihood that they are related to motion or reflect background stochastic fluctuation. In all plots, these 1568 points are opaque. For completeness, we also include the remaining 1556 detected peaks that were discarded. These points are depicted as gray and transparent. (*a*) Definition of several quantities of interest. (*b*) Peak height for three peak types. (*c*) Trough-to-trough durations for three peaks. (*d*) Maximum velocity for three peaks. (*e*) Relationship between peak height and duration. (*f*) Relationship between velocity and duration. (*g*) Relationship between peak height and velocity. (*h*) Mean RMS trough-to-trough curves for co-fluctuation peaks. The inset depicts analogous data but for mean slope, rather than RMS. In this panel, color indicates duration.

**FIG. S2.**
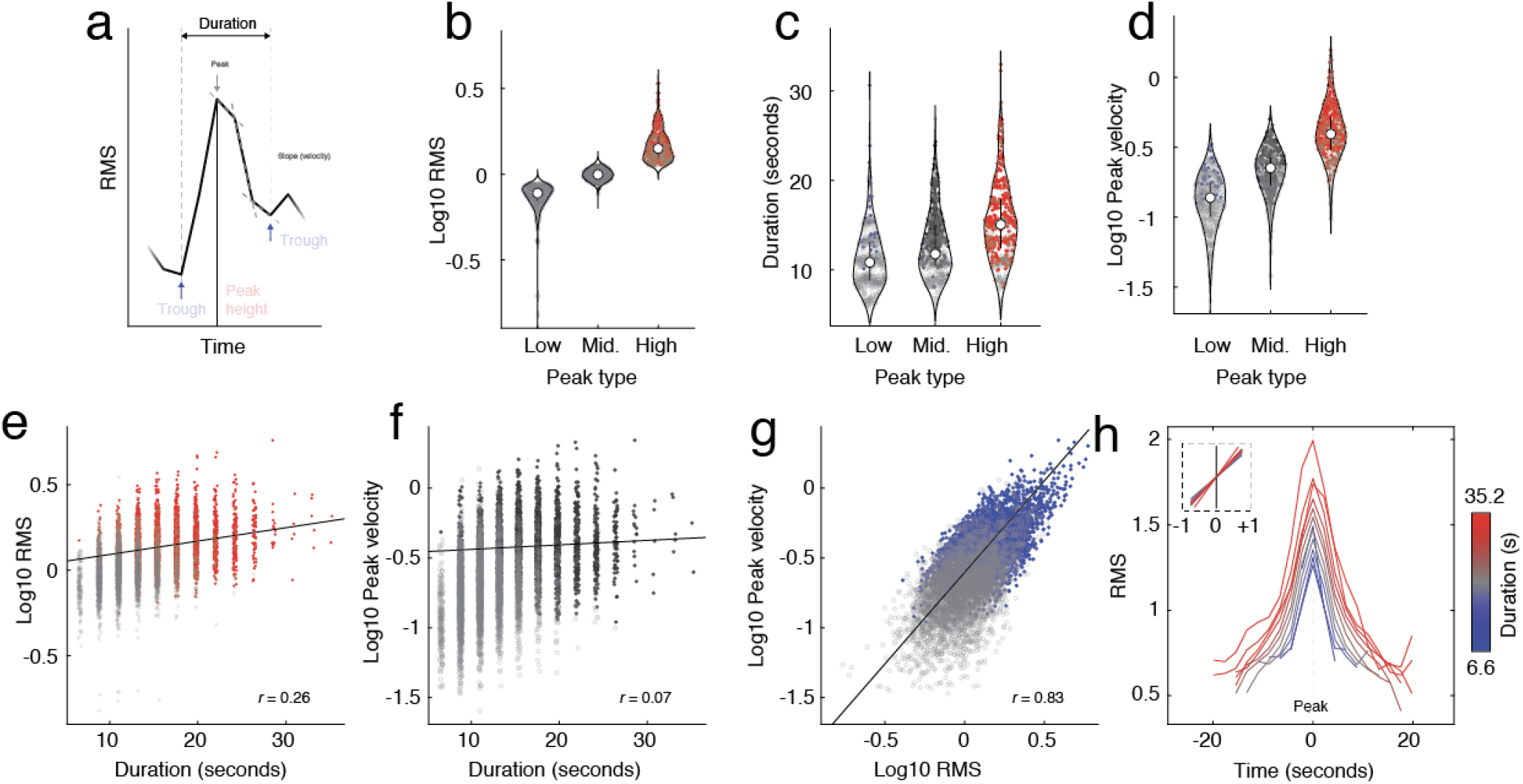
Characterizing peak co-fluctuations in Midnight Scan Club data. In the main text, we characterized peak co-fluctuation using data from the MyConnectome project. Here, we repeated this analysis using data from the Midnight Scan Club after pooling together data from eight subjects (MSC08 and MSC09 were excluded due to data quality issues). For every peak, we calculated its amplitude (RMS) and duration. Using a procedure developed in our previous work, also classified each peak as a high-, middle-, or low-amplitude frame. (*a*) Definition of several quantities of interest. (*b*) Peak height for three peak types. (*c*) Trough-to-trough durations for three peaks. (*d*) Maximum velocity for three peaks. (*e*) Relationship between peak height and duration. (*f*) Relationship between velocity and duration. (*g*) Relationship between peak height and velocity. (*h*) Typical RMS trough-to-trough curves for co-fluctuation peaks. In this panel, color indicates duration.

**FIG. S3.**
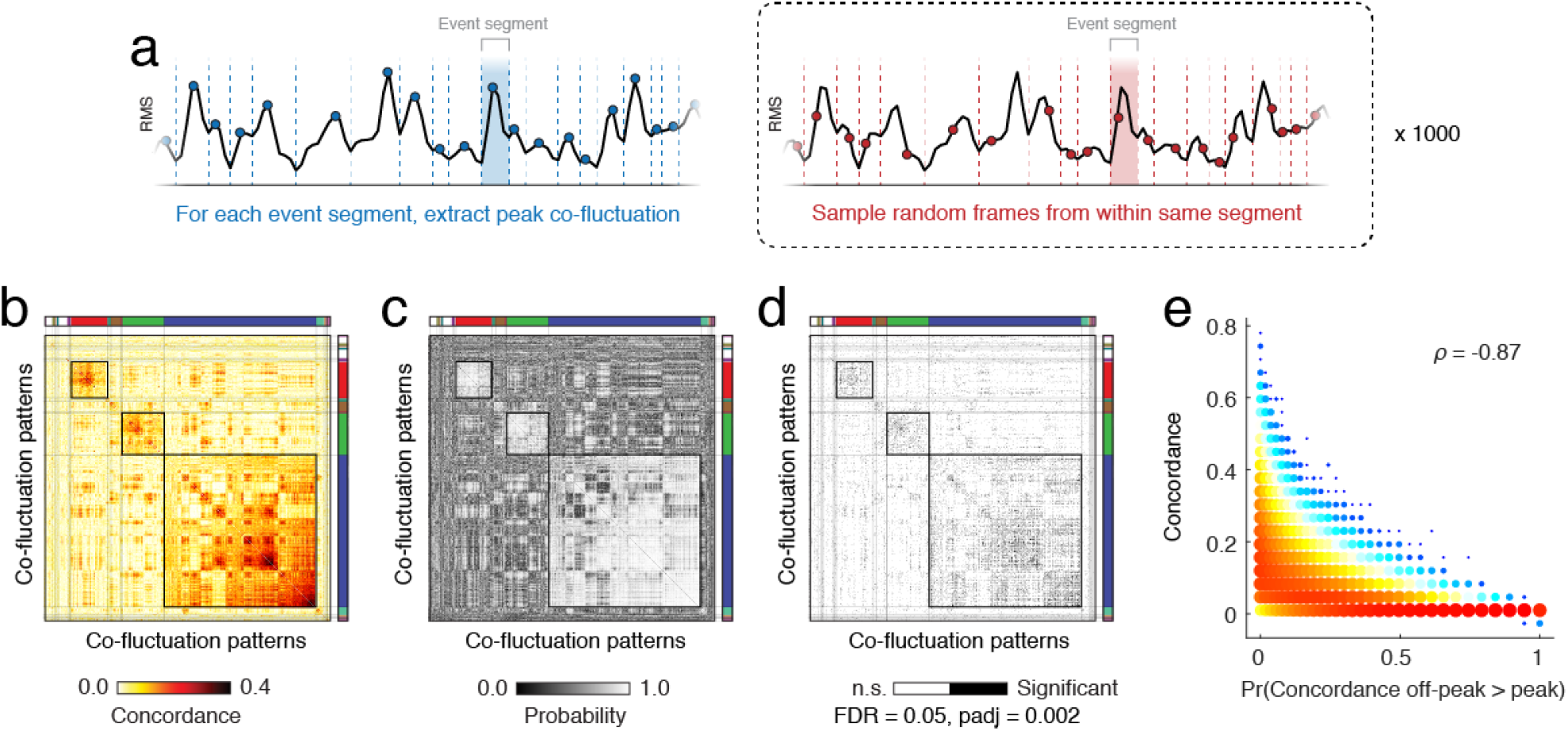
Comparing peak and off-peak co-fluctuation patterns. In the main text, we examine peak co-fluctuation patterns. Here, we compare concordance matrices estimated from peak co-fluctuations against those estimated using patterns taken from within the same event segment but at off-peak frames (see *a* for a schematic of this null model). (*b*) The observed concordance matrix. (*c*) Out of 1000 random samples of off-peak frames, we calculated the fraction of those samples in which the similarity of elements in the null concordance matrix were greater than or equal to those of the observed matrix. Small p-values indicate event segments whose peak-peak concordance was significantly greater than the concordance of random-samples of off-peaks. (*d*) Black cells in this matrix indicate those pairs of segments that survive multiple comparisons corrections (false discovery rate fixed at *q* = 0.05 and the p-value adjusted to *p_adj_* = 0.002. (*e*) Scatterplot of p-values versus observed concordance values. Note that concordant pairs of co-fluctuations tend to have small p-values.

**FIG. S4.**
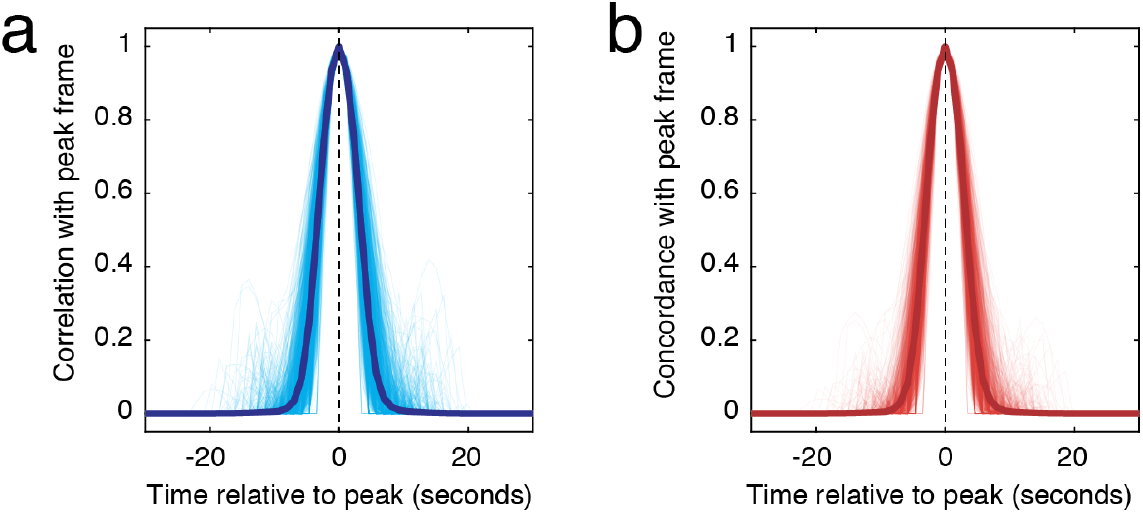
Similarity of co-fluctuation patterns to peak. We analyzed peak co-fluctuation patterns in the main text. Here, we show the similarity of nearby frames in the same segment to those peak patterns. (*a*) Similarity as measured with Pearson’s correlation coefficient. (*b*) Similarity as measured by Lin’s concordance.

**FIG. S5.**
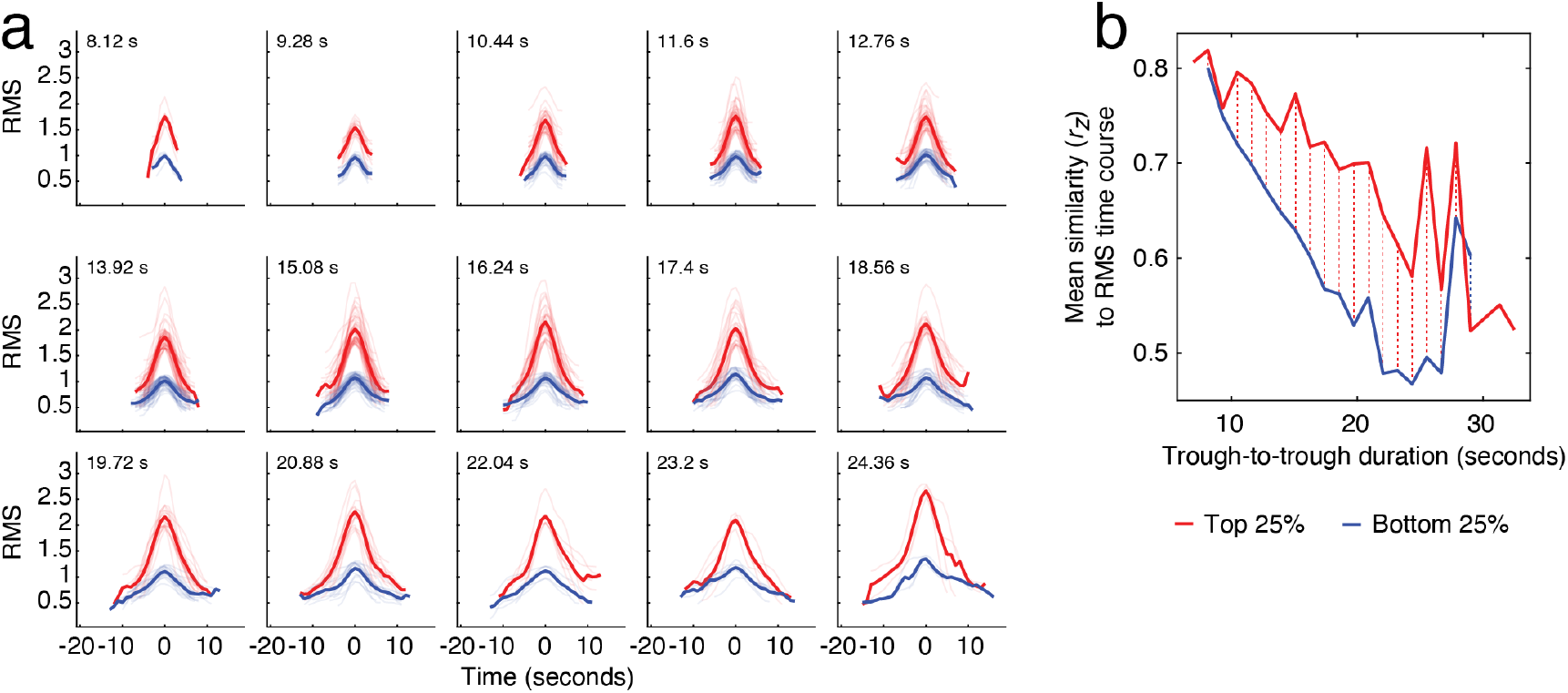
Top and bottom RMS quartiles by duration. In the main text we described a correlation between duration and amplitude of peak co-fluctulations. There was, however, considerable variance around that best linear fit between those variables. (*a*) Here, for each duration (in units of TRs) we show the top (red) and bottom (blue) trough-to-trough curves, ranked by RMS. (*b*) We returned to the edge time series and calculated the correlation of each edge time series with the corresponding trough-to-trough RMS curve. We found that the mean correlation over all edges was stronger for the top 25% than for the bottom 25%, suggesting that even after controlling for duration, there is variability in the “diffusivity” of the trough-to-trough co-fluctuation, with higher amplitude co-fluctuations corresponding to tighter and more cohesive fluctuations than lower-amplitude fluctuations of identical duration.

**FIG. S6.**
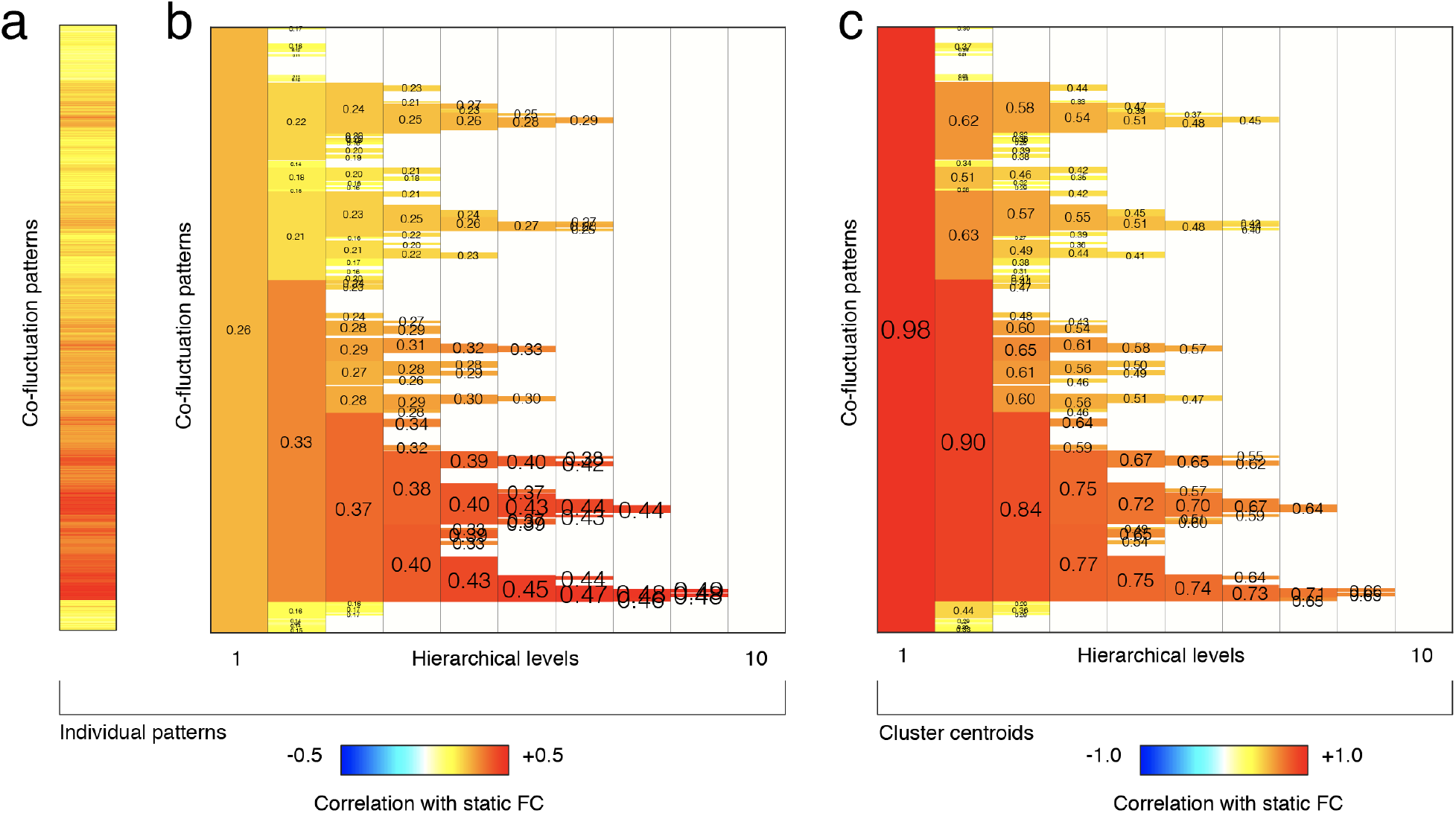
Correlation of cluster centroids and individual patterns with static FC. In Fig. 3j we showed that larger clusters and their centroids were more strongly correlated with FC. Here, we repeat this analysis showing (*a*) the correlation of individual co-fluctuation patterns with FC, (*b*) the mean across those pattern level correlations averaged by cluster, and (*c*) the correlation of cluster centroids with FC.

**FIG. S7.**
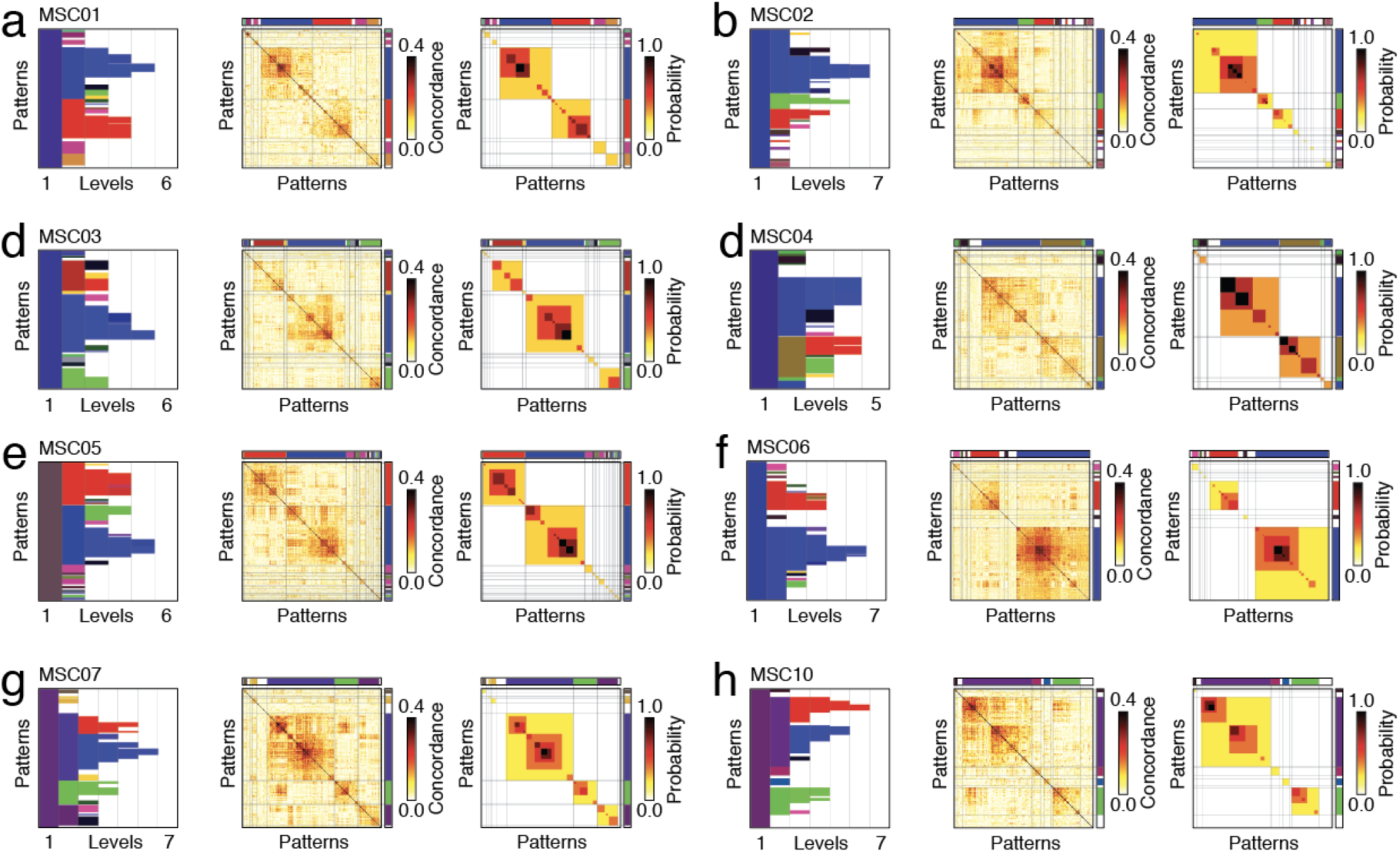
Hierarchical clusters, concondance, and co-assignment matrices. Here, we show results of the hierarchical clustering algorithm applied to data from the Midnight Scan Club. Panels *a-h* show data from MSC01-MSC07 and MSC10. Each panel includes three sub-panels. From left to right: hierarchical cluster labels for each co-fluctuation pattern; concordance matrix ordered by clusters; co-assignment matrix ordered by clusters.

**FIG. S8.**
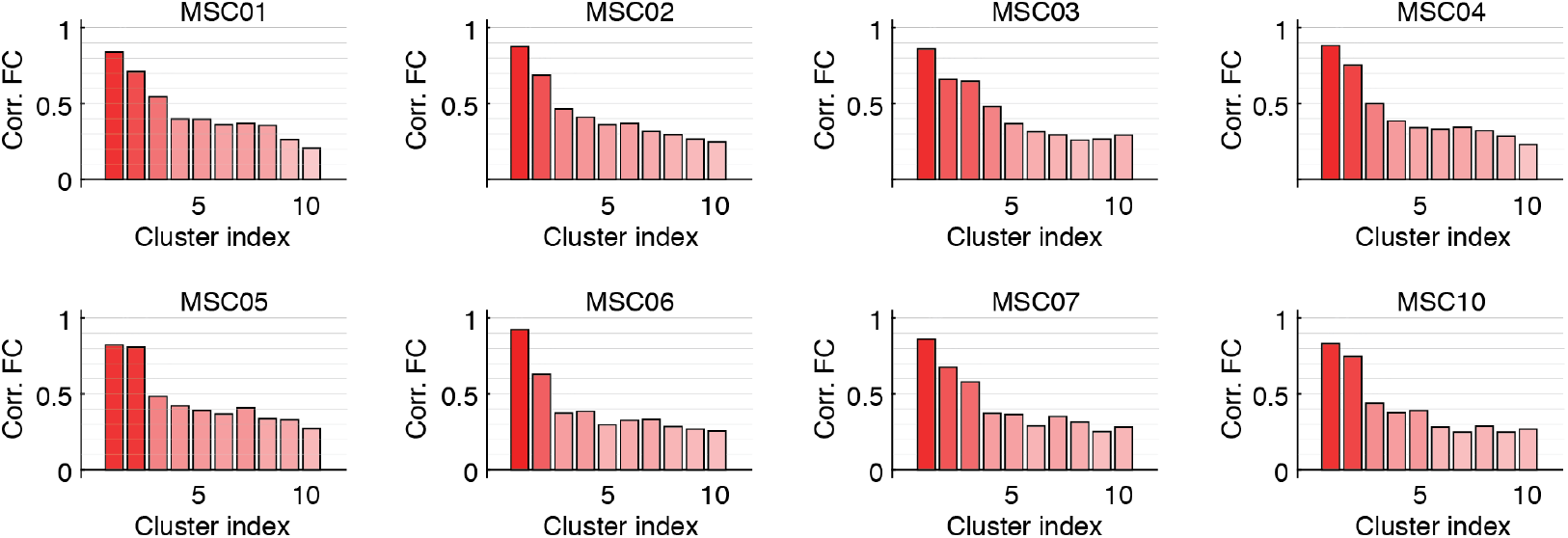
Correlation of cluster centroids with static FC. In Fig. 3j we showed that larger clusters were more strongly correlated with FC. Here, we repeat this analysis using data from the Midnight Scan Club. Each panel correspond to a different subject.

**FIG. S9.**
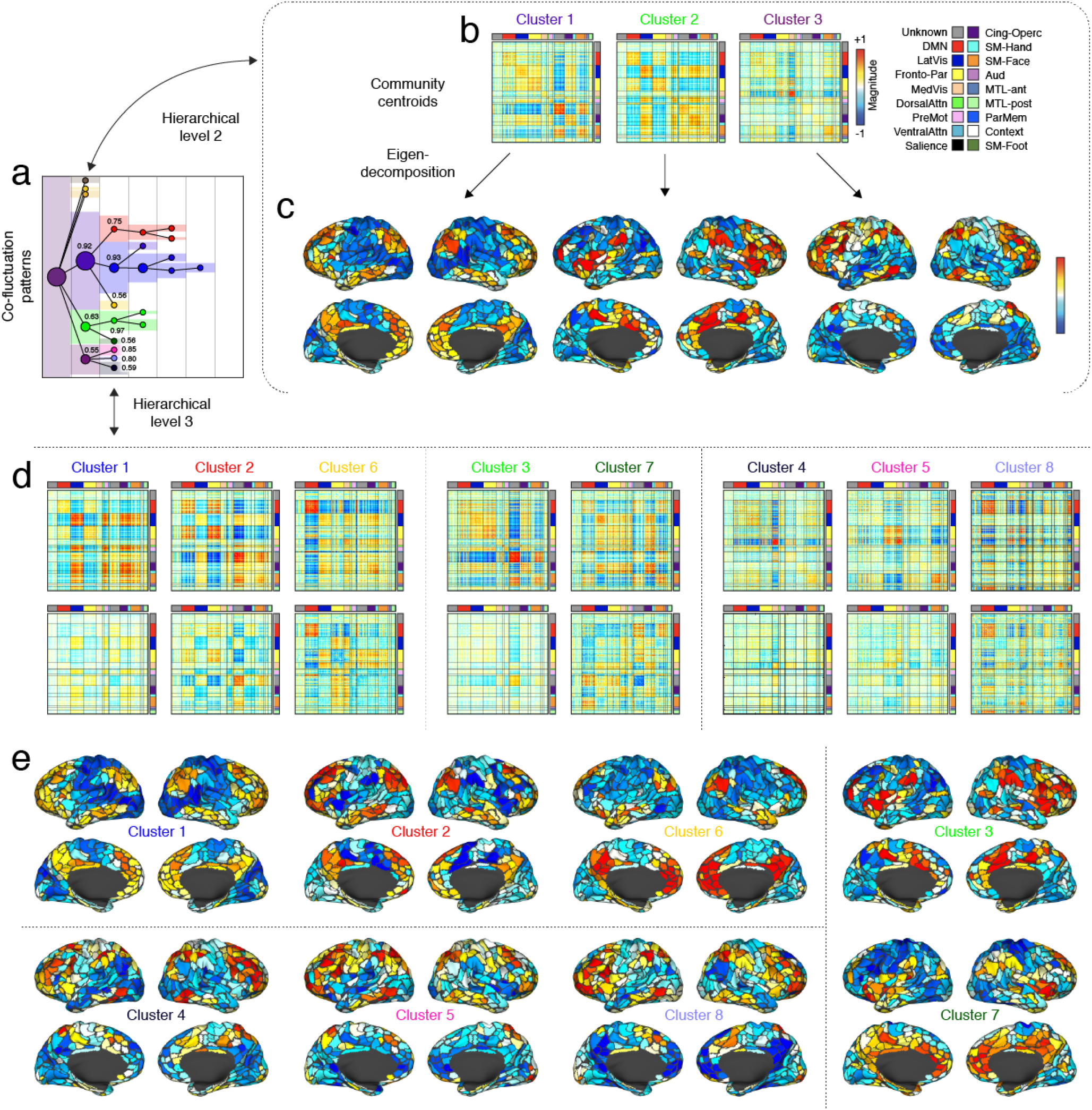
Sub-divisions of clusters in MSC data: An example using MSC07. In the main text we demonstrated that co-fluctuation patterns could be organized hierarchically. In the supplement, we applied the hierarchical clustering algorithm to all MSC participants. For illustrative purposes, we show here the results for subject MSC07. (*a*) Hierarchical cluster assignment with dendrogram overlaid. (*b*) We show the three largest cluster centroids at hierarchical level 2. (*c*) The leading eigenvector for each centroid. (*d*) We show decompositions of clusters 2, 1, and 3 into smaller and more distinct sub-clusters. (e) Again, we show the leading eigenvectors for each sub-cluster.

**FIG. S10.**
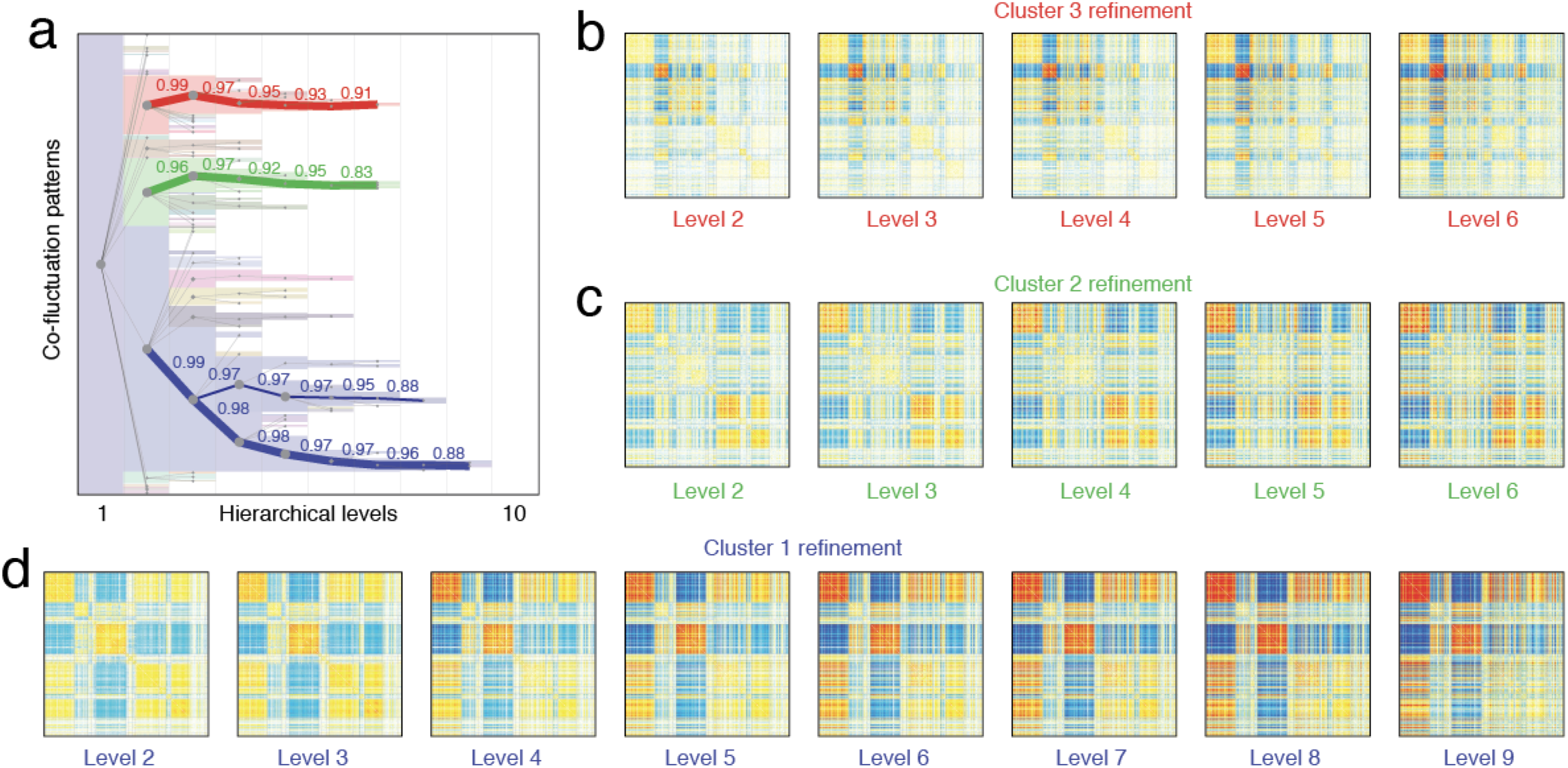
Persistence and refinement of coarse clusters across hierarchical levels. In Fig. 3 and Fig. 4 we showed coarse clusters and their hierarchical divisions. We note that the coarse clusters, although they get sub-divided, are also refined across hierarchical levels. That is, strong co-fluctuations get stronger (positive and negative) but the overall pattern persists. Here, we highlight the persistence of the three large clusters identified in Fig. 3. The correlation values shown in *a* correspond to the correlation of each child centroid with its immediate parent. Note that an alternative possibility was that, as clusters sub-divide, the children partitions decompose their parents so that the correspondence is not as strong.

**FIG. S11.**
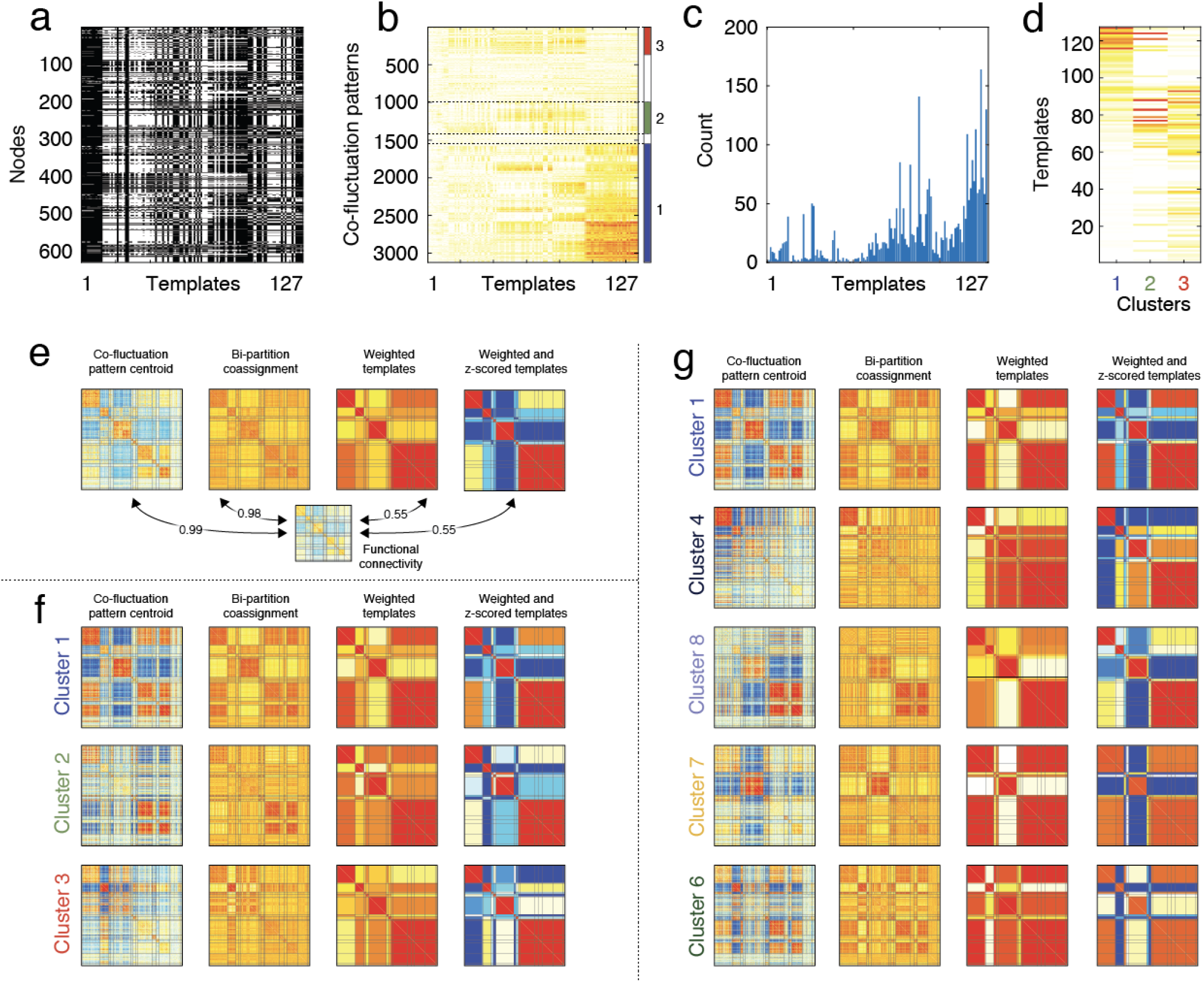
Comparison with templates and bipartitions. (*a*) System templates. Given 14 systems, there are exactly 127 unique templates. (*b*) Normalized mutual information of each co-fluctuation pattern with each template. Patterns are ordered by community. Note: Here we include all co-fluctuation patterns, including those with low prominence and without exclusion based on proximity to another peak. (*c*) For each pattern, we identified the index corresponding to the maximum NMI. Here, we show the histogram. (*d*) We then averaged NMI values within each of the three large clusters at hierarchical level 2 to reveal distinct NMI profiles. (*e*) We then compared different reconstructions of FC to the observed FC matrix. These included the mean co-fluctuation pattern (averaged over all patterns), the co-assignment matrix of observed bipartition, the co-assignment matrix of the best-matched system-templates, and the z-scored version of the best-matched templates. We repeated this analysis for the largest clusters detected at hierarchical level 2 (*f*) and for subdivisions of the largest cluster at that level (*g*). In all cases, we find evidence that templates capture the specificity of divisions among co-fluctuations originally identified using the hierarchical clustering algorithm.

**FIG. S12.**
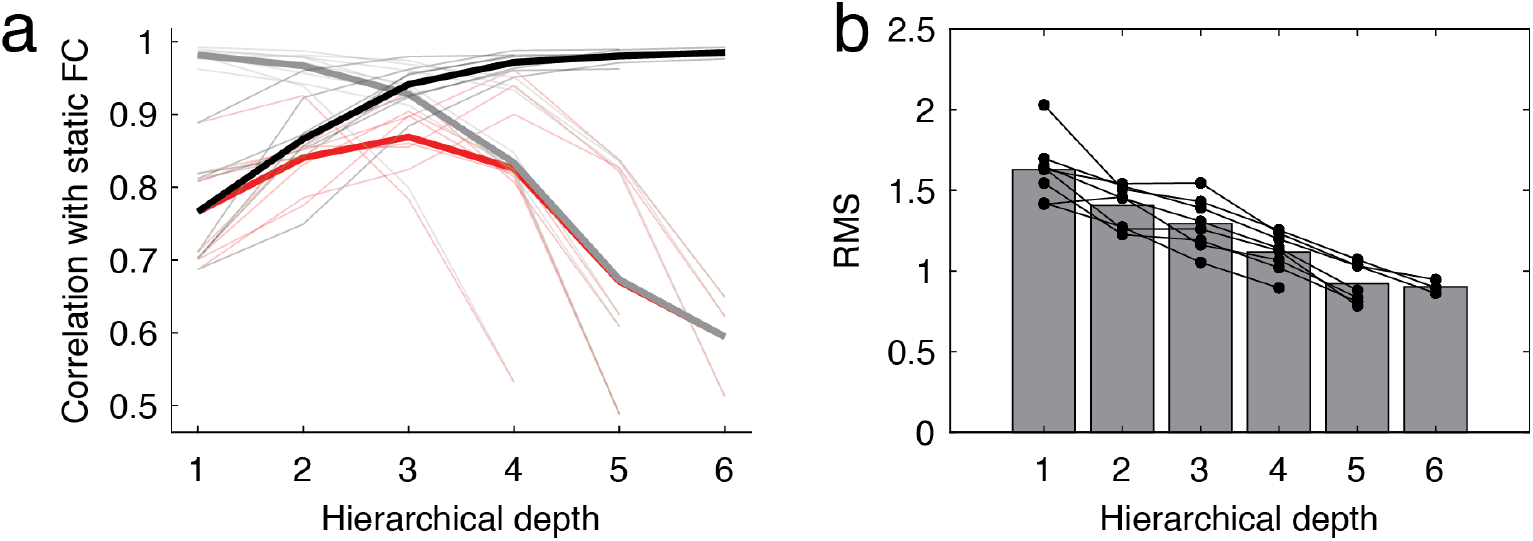
Linking different hierarchical levels to FC using MSC data. In the main text we show that the correspondence with FC peaks at an intermediate hierarchical level. Here, we recapitulate that analysis using data from the individual subjects in the MSC dataset. (*a*) Correlation with FC at different hierarchical levels. Thick lines indicate subject-averages, thin lines indicate data from individual subjects. (*b*) Mean RMS of co-fluctuation patterns at different hierarchical levels.

## Footnotes

1 We discovered later that this method had been reported at least once before, but had been applied in a narrow context [47].

## References

[1] Danielle S Bassett and Olaf Sporns, “Network neuroscience,” Nature neuroscience 20, 353–364 (2017).

[2] Andrew T Reid, Drew B Headley, Ravi D Mill, Ruben Sanchez-Romero, Lucina Q Uddin, Daniele Marinazzo, Daniel J Lurie, Pedro A Valdés-Sosa, Stephen José Hanson, Bharat B Biswal, et al., “Advancing functional connectivity research from association to causation,” Nature neuroscience 22, 1751–1760 (2019).

[3] Farnaz Zamani Esfahlani, Youngheun Jo, Joshua Faskowitz, Lisa Byrge, Daniel Kennedy, Olaf Sporns, and Richard Betzel, “High-amplitude co-fluctuations in cortical activity drive functional connectivity,” Proceedings of the National Academy of Sciences (2020).

[4] Joshua Faskowitz, Farnaz Zamani Esfahlani, Youngheun Jo, Olaf Sporns, and Richard F Betzel, “Edge-centric functional network representations of human cerebral cortex reveal overlapping system-level architecture,” Nature neuroscience 23, 1644–1654 (2020).

[5] Olaf Sporns, Joshua Faskowitz, Andreia Sofia Teixeira, Sarah A Cutts, and Richard F Betzel, “Dynamic expression of brain functional systems disclosed by fine-scale analysis of edge time series,” Network Neuroscience 5, 405–433 (2021).

[6] Sarah Greenwell, Joshua Faskowitz, Laura Pritschet, Tyler Santander, Emily G Jacobs, and Richard F Betzel, “High-amplitude network co-fluctuations linked to variation in hormone concentrations over menstrual cycle,” bioRxiv (2021).

[7] Richard Betzel, Sarah Cutts, Sarah Greenwell, and Olaf Sporns, “Individualized event structure drives individual differences in whole-brain functional connectivity,” bioRxiv (2021).

[8] Farnaz Zamani Esfahlani, Lisa Byrge, Jacob Tanner, Olaf Sporns, Daniel Kennedy, and Richard Betzel, “Edge-centric analysis of time-varying functional brain networks with applications in autism spectrum disorder,” bioRxiv (2021).

[9] Youngheun Jo, Joshua Faskowitz, Farnaz Zamani Esfahlani, Olaf Sporns, and Richard F Betzel, “Subject identification using edge-centric functional connectivity,” bioRxiv (2020).

[10] Enzo Tagliazucchi, Pablo Balenzuela, Daniel Fraiman, and Dante R Chialvo, “Criticality in large-scale brain fmri dynamics unveiled by a novel point process analysis,” Frontiers in physiology 3, 15 (2012).

[11] Xiao Liu, Catie Chang, and Jeff H Duyn, “Decomposition of spontaneous brain activity into distinct fmri co-activation patterns,” Frontiers in systems neuroscience 7, 101 (2013).

[12] Thomas W Allan, Susan T Francis, Cesar Caballero-Gaudes, Peter G Morris, Elizabeth B Liddle, Peter F Liddle, Matthew J Brookes, and Penny A Gowland, “Functional connectivity in mri is driven by spontaneous bold events,” PloS one 10, e0124577 (2015).

[13] I Cifre, M Zarepour, SG Horovitz, SA Cannas, and DR Chialvo, “Further results on why a point process is effective for estimating correlation between brain regions,” Papers in Physics 12, 120003–120003 (2020).

[14] Natalia Petridou, César Caballero Gaudes, Ian L Dryden, Susan T Francis, and Penny A Gowland, “Periods of rest in fmri contain individual spontaneous events which are related to slowly fluctuating spontaneous activity,” Human brain mapping 34, 1319–1329 (2013).

[15] Timothy O Laumann, Evan M Gordon, Babatunde Adeyemo, Abraham Z Snyder, Sung Jun Joo, Mei-Yen Chen, Adrian W Gilmore, Kathleen B McDermott, Steven M Nelson, Nico UF Dosenbach, et al., “Functional system and areal organization of a highly sampled individual human brain,” Neuron 87, 657–670 (2015).

[16] Russell A Poldrack, “Precision neuroscience: Dense sampling of individual brains,” Neuron 95, 727–729 (2017).

[17] Evan M Gordon, Timothy O Laumann, Adrian W Gilmore, Dillan J Newbold, Deanna J Greene, Jeffrey J Berg, Mario Ortega, Catherine Hoyt-Drazen, Caterina Gratton, Haoxin Sun, et al., “Precision functional mapping of individual human brains,” Neuron 95, 791–807 (2017).

[18] Caterina Gratton, Timothy O Laumann, Ashley N Nielsen, Deanna J Greene, Evan M Gordon, Adrian W Gilmore, Steven M Nelson, Rebecca S Coalson, Abraham Z Snyder, Bradley L Schlaggar, et al., “Functional brain networks are dominated by stable group and individual factors, not cognitive or daily variation,” Neuron 98, 439–452 (2018).

[19] Mark EJ Newman and Michelle Girvan, “Finding and evaluating community structure in networks,” Physical review E 69, 026113 (2004).

[20] Vincent D Blondel, Jean-Loup Guillaume, Renaud Lambiotte, and Etienne Lefebvre, “Fast unfolding of communities in large networks,” Journal of statistical mechanics: theory and experiment 2008, P10008 (2008).

[21] I Lawrence and Kuei Lin, “A concordance correlation coefficient to evaluate reproducibility,” Biometrics, 255–268 (1989).

[22] Maxwell Shinn, Amber Hu, Laurel Turner, Stephanie Noble, Sophie Achard, Alan Anticevic, Dustin Scheinost, R Todd Constable, Daeyeol Lee, Edward T Bullmore, et al., “Spatial and temporal autocorrelation weave human brain networks,” bioRxiv (2021).

[23] Richard F Betzel and Danielle S Bassett, “Multi-scale brain networks,” Neuroimage 160, 73–83 (2017).

[24] David Meunier, Renaud Lambiotte, Alex Fornito, Karen Ersche, and Edward T Bullmore, “Hierarchical modularity in human brain functional networks,” Frontiers in neuroinformatics 3, 37 (2009).

[25] Danielle S Bassett and Edward T Bullmore, “Human brain networks in health and disease,” Current opinion in neurology 22, 340 (2009).

[26] CJ van Stam and ECW Van Straaten, “The organization of physiological brain networks,” Clinical neurophysiology 123, 1067–1087 (2012).

[27] Changsong Zhou, Lucia Zemanová, Gorka Zamora, Claus C Hilgetag, and Jürgen Kurths, “Hierarchical organization unveiled by functional connectivity in complex brain networks,” Physical review letters 97, 238103 (2006).

[28] Arian Ashourvan, Qawi K Telesford, Timothy Verstynen, Jean M Vettel, and Danielle S Bassett, “Multiscale detection of hierarchical community architecture in structural and functional brain networks,” Plos one 14, e0215520 (2019).

[29] Claus C Hilgetag and Alexandros Goulas, “‘hierarchy’in the organization of brain networks,” Philosophical Transactions of the Royal Society B 375, 20190319 (2020).

[30] Daniel J Lurie, Daniel Kessler, Danielle S Bassett, Richard F Betzel, Michael Breakspear, Shella Kheilholz, Aaron Kucyi, Raphael Liégeois, Martin A Lindquist, Anthony Randal McIntosh, et al., “Questions and controversies in the study of time-varying functional connectivity in resting fmri,” Network Neuroscience 4, 30–69 (2020).

[31] Sepideh Sadaghiani, Matthew J Brookes, and Sylvain Baillet, “Connectomics of human electrophysiology,” NeuroImage 247, 118788 (2022).

[32] Marika Strindberg, Peter Fransson, Joana Cabral, and Ulrika Ådén, “Spatiotemporally flexible subnetworks reveal the quasi-cyclic nature of integration and segregation in the human brain,” NeuroImage 239, 118287 (2021).

[33] Karolina Finc, Kamil Bonna, Xiaosong He, David M Lydon-Staley, Simone Kuhn, Włodzislaw Duch, and Danielle S Bassett, “Dynamic reconfiguration of functional brain networks during working memory training,” Nature communications 11, 1–15 (2020).

[34] Danielle S Bassett, Nicholas F Wymbs, Mason A Porter, Peter J Mucha, Jean M Carlson, and Scott T Grafton, “Dynamic reconfiguration of human brain networks during learning,” Proceedings of the National Academy of Sciences 108, 7641–7646 (2011).

[35] Elena A Allen, Eswar Damaraju, Sergey M Plis, Erik B Erhardt, Tom Eichele, and Vince D Calhoun, “Tracking whole-brain connectivity dynamics in the resting state,” Cerebral cortex 24, 663–676 (2014).

[36] Nora Leonardi, William R Shirer, Michael D Greicius, and Dimitri Van De Ville, “Disentangling dynamic networks: Separated and joint expressions of functional connectivity patterns in time,” Human brain mapping 35, 5984–5995 (2014).

[37] Timothy O Laumann, Abraham Z Snyder, Anish Mitra, Evan M Gordon, Caterina Gratton, Babatunde Adeyemo, Adrian W Gilmore, Steven M Nelson, Jeff J Berg, Deanna J Greene, et al., “On the stability of bold fmri correlations,” Cerebral cortex 27, 4719–4732 (2017).

[38] Raphael Liegeois, Timothy O Laumann, Abraham Z Snyder, Juan Zhou, and BT Thomas Yeo, “Interpreting temporal fluctuations in resting-state functional connectivity mri,” Neuroimage 163, 437–455 (2017).

[39] Eli J Cornblath, Arian Ashourvan, Jason Z Kim, Richard F Betzel, Rastko Ciric, Azeez Adebimpe, Graham L Baum, Xiaosong He, Kosha Ruparel, Tyler M Moore, et al., “Temporal sequences of brain activity at rest are constrained by white matter structure and modulated by cognitive demands,” Communications biology 3, 1–12 (2020).

[40] Barnaly Rashid, Eswar Damaraju, Godfrey D Pearlson, and Vince D Calhoun, “Dynamic connectivity states estimated from resting fmri identify differences among schizophrenia, bipolar disorder, and healthy control subjects,” Frontiers in human neuroscience 8, 897 (2014).

[41] John D Medaglia, Theodore D Satterthwaite, Apoorva Kelkar, Rastko Ciric, Tyler M Moore, Kosha Ruparel, Ruben C Gur, Raquel E Gur, and Danielle S Bassett, “Brain state expression and transitions are related to complex executive cognition in normative neurodevelopment,” Neuroimage 166, 293–306 (2018).

[42] Javier Gonzalez-Castillo, Colin W Hoy, Daniel A Handwerker, Meghan E Robinson, Laura C Buchanan, Ziad S Saad, and Peter A Bandettini, “Tracking ongoing cognition in individuals using brief, whole–brain functional connectivity patterns,” Proceedings of the National Academy of Sciences 112, 8762–8767 (2015).

[43] Eleonora Fiorenzato, Antonio P Strafella, Jinhee Kim, Roberta Schifano, Luca Weis, Angelo Antonini, and Roberta Biundo, “Dynamic functional connectivity changes associated with dementia in parkinson’s disease,” Brain 142, 2860–2872 (2019).

[44] Rikkert Hindriks, Mohit H Adhikari, Yusuke Murayama, Marco Ganzetti, Dante Mantini, Nikos K Logothetis, and Gustavo Deco, “Can sliding-window correlations reveal dynamic functional connectivity in resting-state fmri?” Neuroimage 127, 242–256 (2016).

[45] Nora Leonardi and Dimitri Van De Ville, “On spurious and real fluctuations of dynamic functional connectivity during rest,” Neuroimage 104, 430–436 (2015).

[46] Andrew Zalesky and Michael Breakspear, “Towards a statistical test for functional connectivity dynamics,” Neuroimage 114, 466–470 (2015).

[47] Erik SB van Oort, Maarten Mennes, Tobias Navarro Schröder, Vinod J Kumar, Nestor I Zaragoza Jimenez, Wolfgang Grodd, Christian F Doeller, and Christian F Beckmann, “Human brain parcellation using time courses of instantaneous connectivity,” arXiv preprint arXiv:1609.04636 (2016).

[48] Robyn L Miller, Maziar Yaesoubi, Jessica A Turner, Daniel Mathalon, Adrian Preda, Godfrey Pearlson, Tulay Adali, and Vince D Calhoun, “Higher dimensional meta-state analysis reveals reduced resting fmri connectivity dynamism in schizophrenia patients,” PloS one 11, e0149849 (2016).

[49] Gaëlle Doucet, Mikaël Naveau, Laurent Petit, Nicolas Delcroix, Laure Zago, Fabrice Crivello, Gael Jobard, Nathalie Tzourio-Mazoyer, Bernard Mazoyer, Emmanuel Mellet, et al., “Brain activity at rest: a multiscale hierarchical functional organization,” Journal of neurophysiology 105, 2753–2763 (2011).

[50] Stephen M Smith, Karla L Miller, Steen Moeller, Junqian Xu, Edward J Auerbach, Mark W Woolrich, Christian F Beckmann, Mark Jenkinson, Jesper Andersson, Matthew F Glasser, et al., “Temporally-independent functional modes of spontaneous brain activity,” Proceedings of the National Academy of Sciences 109, 3131–3136 (2012).

[51] Diego Vidaurre, Stephen M Smith, and Mark W Woolrich, “Brain network dynamics are hierarchically organized in time,” Proceedings of the National Academy of Sciences 114, 12827–12832 (2017).

[52] Michael D Fox, Abraham Z Snyder, Justin L Vincent, Maurizio Corbetta, David C Van Essen, and Marcus E Raichle, “The human brain is intrinsically organized into dynamic, anticorrelated functional networks,” Proceedings of the National Academy of Sciences 102, 9673–9678 (2005).

[53] Yulia Golland, Polina Golland, Shlomo Bentin, and Rafael Malach, “Data-driven clustering reveals a fundamental subdivision of the human cortex into two global systems,” Neuropsychologia 46, 540–553 (2008).

[54] Taylor Bolt, Jason S Nomi, Danilo Bzdok, Catie Chang, BT Thomas Yeo, Lucina Q Uddin, and Shella D Keilholz, “Large-scale intrinsic functional brain organization emerges from three canonical spatiotemporal patterns,” bioRxiv (2021).

[55] Daniel S Margulies, Satrajit S Ghosh, Alexandros Goulas, Marcel Falkiewicz, Julia M Huntenburg, Georg Langs, Gleb Bezgin, Simon B Eickhoff, F Xavier Castellanos, Michael Petrides, et al., “Situating the default-mode network along a principal gradient of macroscale cortical organization,” Proceedings of the National Academy of Sciences 113, 12574–12579 (2016).

[56] M-Marsel Mesulam, “From sensation to cognition.” Brain: a journal of neurology 121, 1013–1052 (1998).

[57] Andrea Avena-Koenigsberger, Bratislav Misic, and Olaf Sporns, “Communication dynamics in complex brain networks,” Nature Reviews Neuroscience 19, 17 (2018).

[58] Z Ladwig, BA Seitzman, A Dworetsky, Y Yu, B Adeyemo, DM Smith, SE Petersen, and C Gratton, “Bold cofluctuation ‘events’ are predicted from static functional connectivity,” (2022).

[59] Stephanie Noble, Marisa N Spann, Fuyuze Tokoglu, Xilin Shen, R Todd Constable, and Dustin Scheinost, “Influences on the test-retest reliability of functional connectivity mri and its relationship with behavioral utility,” Cerebral cortex 27, 5415–5429 (2017).

[60] Leonardo Novelli and Adeel Razi, “A mathematical perspective on edge-centric functional connectivity,” arXiv preprint arXiv:2106.10631 (2021).

[61] Teppei Matsui, Trung Quang Pham, Koji Jimura, and Junichi Chikazoe, “On co-activation pattern analysis and non-stationarity of resting brain activity,” NeuroImage, 118904 (2022).

[62] Petra Ritter, Michael Schirner, Anthony R McIntosh, and Viktor K Jirsa, “The virtual brain integrates computational modeling and multimodal neuroimaging,” Brain connectivity 3, 121–145 (2013).

[63] Paula Sanz-Leon, Stuart A Knock, Andreas Spiegler, and Viktor K Jirsa, “Mathematical framework for large-scale brain network modeling in the virtual brain,” Neuroimage 111, 385–430 (2015).

[64] Joana Cabral, Morten L Kringelbach, and Gustavo Deco, “Functional connectivity dynamically evolves on multiple time-scales over a static structural connectome: Models and mechanisms,” Neuroimage 160, 84–96 (2017).

[65] Raphael Liégeois, BT Thomas Yeo, and Dimitri Van De Ville, “Interpreting null models of resting-state functional mri dynamics: not throwing the model out with the hypothesis,” Neuroimage 243, 118518 (2021).

[66] Maria Pope, Makoto Fukushima, Richard Betzel, and Olaf Sporns, “Modular origins of high-amplitude co-fluctuations in fine-scale functional connectivity dynamics,” bioRxiv (2021).

[67] Megan M Sperry, Sonia Kartha, Eric J Granquist, and Beth A Winkelstein, “Inter-subject fdg pet brain networks exhibit multi-scale community structure with different normalization techniques,” Annals of biomedical engineering 46, 1001–1012 (2018).

[68] Evan M Gordon, Timothy O Laumann, Scott Marek, Ryan V Raut, Caterina Gratton, Dillan J Newbold, Deanna J Greene, Rebecca S Coalson, Abraham Z Snyder, Bradley L Schlaggar, et al., “Default-mode network streams for coupling to language and control systems,” Proceedings of the National Academy of Sciences 117, 17308–17319 (2020).

[69] Roger Guimera, Marta Sales-Pardo, and Luis A NunesAmaral, “Modularity from fluctuations in random graphs and complex networks,” Physical Review E 70, 025101 (2004).

[70] Lucas GS Jeub, Olaf Sporns, and Santo Fortunato, “Multiresolution consensus clustering in networks,” Scientific reports 8, 1–16 (2018).

[71] Anders M Dale, Bruce Fischl, and Martin I Sereno, “Cortical surface-based analysis: I. segmentation and surface reconstruction,” Neuroimage 9, 179–194 (1999).

[72] Matthew F Glasser, Stamatios N Sotiropoulos, J Anthony Wilson, Timothy S Coalson, Bruce Fischl, Jesper L Andersson, Junqian Xu, Saad Jbabdi, Matthew Webster, Jonathan R Polimeni, et al., “The minimal preprocessing pipelines for the human connectome project,” Neuroimage 80, 105–124 (2013).

[73] Jean Talairach, “Co-planar stereotaxic atlas of the human brain-3-dimensional proportional system,” An approach to cerebral imaging (1988).

[74] Stephen M Smith, Mark Jenkinson, Mark W Woolrich, Christian F Beckmann, Timothy EJ Behrens, Heidi Johansen-Berg, Peter R Bannister, Marilena De Luca, Ivana Drobnjak, David E Flitney, et al., “Advances in functional and structural mr image analysis and implementation as fsl,” Neuroimage 23, S208–S219 (2004).

[75] Jonathan D Power, Anish Mitra, Timothy O Laumann, Abraham Z Snyder, Bradley L Schlaggar, and Steven E Petersen, “Methods to detect, characterize, and remove motion artifact in resting state fmri,” Neuroimage 84, 320–341 (2014).

[76] Rastko Ciric, Daniel H Wolf, Jonathan D Power, David R Roalf, Graham L Baum, Kosha Ruparel, Russell T Shinohara, Mark A Elliott, Simon B Eickhoff, Christos Davatzikos, et al., “Benchmarking of participant-level confound regression strategies for the control of motion artifact in studies of functional connectivity,” Neuroimage 154, 174–187 (2017).

[77] Jonathan D Power, Kelly A Barnes, Abraham Z Snyder, Bradley L Schlaggar, and Steven E Petersen, “Spurious but systematic correlations in functional connectivity mri networks arise from subject motion,” Neuroimage 59, 2142–2154 (2012).

[78] Evan M Gordon, Timothy O Laumann, Babatunde Adeyemo, and Steven E Petersen, “Individual variability of the system-level organization of the human brain,” Cerebral cortex 27, 386–399 (2017).

[79] Russell A Poldrack, Timothy O Laumann, Oluwasanmi Koyejo, Brenda Gregory, Ashleigh Hover, Mei-Yen Chen, Krzysztof J Gorgolewski, Jeffrey Luci, Sung Jun Joo, Ryan L Boyd, et al., “Long-term neural and physiological phenotyping of a single human,” Nature communications 6, 1–15 (2015).

[80] Richard F Betzel, John D Medaglia, Lia Papadopoulos, Graham L Baum, Ruben Gur, Raquel Gur, David Roalf, Theodore D Satterthwaite, and Danielle S Bassett, “The modular organization of human anatomical brain networks: Accounting for the cost of wiring,” Network Neuroscience 1, 42–68 (2017).

